# A structural and dynamic visualization of the interaction between the microtubule-associated protein 7 (MAP7) and microtubules

**DOI:** 10.1101/2023.06.02.543398

**Authors:** Agnes Adler, Mamata Bangera, Salima Bahri, Hugo van Ingen, Carolyn A. Moores, Marc Baldus

## Abstract

Microtubules (MTs) are key components of the eukaryotic cytoskeleton and are essential for intracellular organization, organelle trafficking and mitosis. MT tasks depend on binding and interactions with MT-associated proteins (MAPs). MT-associated protein 7 (MAP7) has the unusual ability of both MT binding and activating kinesin-1-mediated cargo transport along MTs. Additionally, the protein is reported to stabilize MTs with its 112 amino-acid long MT-binding domain (MTBD). Here we investigate the structural basis of the interaction of MAP7 MTBD with the MT lattice. Using a combination of solid and solution-state nuclear magnetic resonance (NMR) spectroscopy with electron microscopy, fluorescence anisotropy and isothermal titration calorimetry, we shed light on the binding mode of MAP7 to MTs at an atomic level. Our results show that a combination of interactions between MAP7 and MT lattice extending beyond a single tubulin dimer and including tubulin C-terminal tails contribute to formation of the MAP7-MT complex.

## Introduction

Mitosis, cell migration, and polarization are dependent on microtubules (MTs). An imbalance in their structure or function is associated with human disorders such as ciliopathies, cancer and neurodegeneration ^1^. These biopolymers are hollow cylinders of α/β-tubulin heterodimers interacting in a “head-to-tail” and side by side fashion ^2,3^. MTs are highly dynamic structures that undergo continuous assembly and disassembly both *in vitro* and *in vivo* in a process termed “dynamic instability” ^4^. This process is driven by hydrolysis of guanosine 5′-triphosphate (GTP) in the MT lattice ^5^. MT-associated proteins (MAPs) interact with MTs, regulate their dynamics, and mediate MT functions in physiological contexts through sophisticated regulation ^6^. The roles and mechanisms of MAPs such as MAP1, MAP2, MAP4, Tau, MAP6, DCX, MAP7 and MAP9 have recently been identified and characterized, particularly in the context of MT-rich neurons ^6,7^. The less well understood MAP7, also known as E-MAP-115 or Ensconsin, is found to bind and stabilize MTs in axons ^8,9^, possibly playing a role in the development of axonal branches. It has been implicated in the growth of metaphase spindles in neural stem cells ^10,11^ and its upregulation has been observed in different forms of cancer ^12–14^. Additionally, MAP7 is a required cofactor for kinesin-1-driven transport along MTs ^15^. Mutations in the gene encoding MAP7 and/or kinesin-1 affect nuclear positioning in myotubes of *Drosophila* embryos as well as mammalian cells leading to muscle defects ^16^.

Shedding light on the MAP7–MT interaction is important for understanding its physiological function and the wider regulation of molecular processes involving the cytoskeleton. The interaction of MAP7 with MTs is mediated by its 112 residues long MT binding domain (MTBD) (residue 59-170) **(Fig. 1a)**. According to solution-state resonance NMR assignments, the MAP7 MTBD exhibits a long α-helix with a short hinge region comprising residues 84-87 ^17^. A recent cryo-electron microscopy (cryo-EM) study of the MAP7-MT complex revealed a 53 residue-long α-helix (residues 87-139) ^18^ bound to the MT (PDBid:7SGS) between the outer protofilament ridge and the inter-protofilament lateral contacts. However, a comprehensive view of the structural organization of the entire MAP7 MTBD on MTs as well as insights into the atomic interactions between MAP7 and the C-terminal tails (CTTs) of α- and β-tubulin, as suggested by previous solid-state NMR experiments ^19^ are currently missing. These CTTs are subject to a variety of post-translational modifications (PTMs) ^20^ affecting the binding of many MAPs to MTs ^18,21,22^, but little is known about the mechanism(s) by which they contribute to MT binding partner interactions. Although cryo-EM has made great progress in unravelling the structure of MTs and of MAP-MT complexes giving insights into their atomic structure and organisation ^6,23^, information about the dynamic segments of MAPs that often contain large intrinsically disordered regions (IDRs) as well as the CTTs is largely missing. NMR has the advantage of not being limited by protein dynamics, making it a suitable tool to study flexible MAP-MT interactions ^24,25^. In addition, the use of magic angle spinning (MAS) solid-state NMR allows for the investigation of large biomolecules ^26,27^. For example, solid-state NMR has been used to study small ligands ^28,29^ as well as MAPs and motor proteins binding to MT ^19,30–34^.

**Fig. 1:**
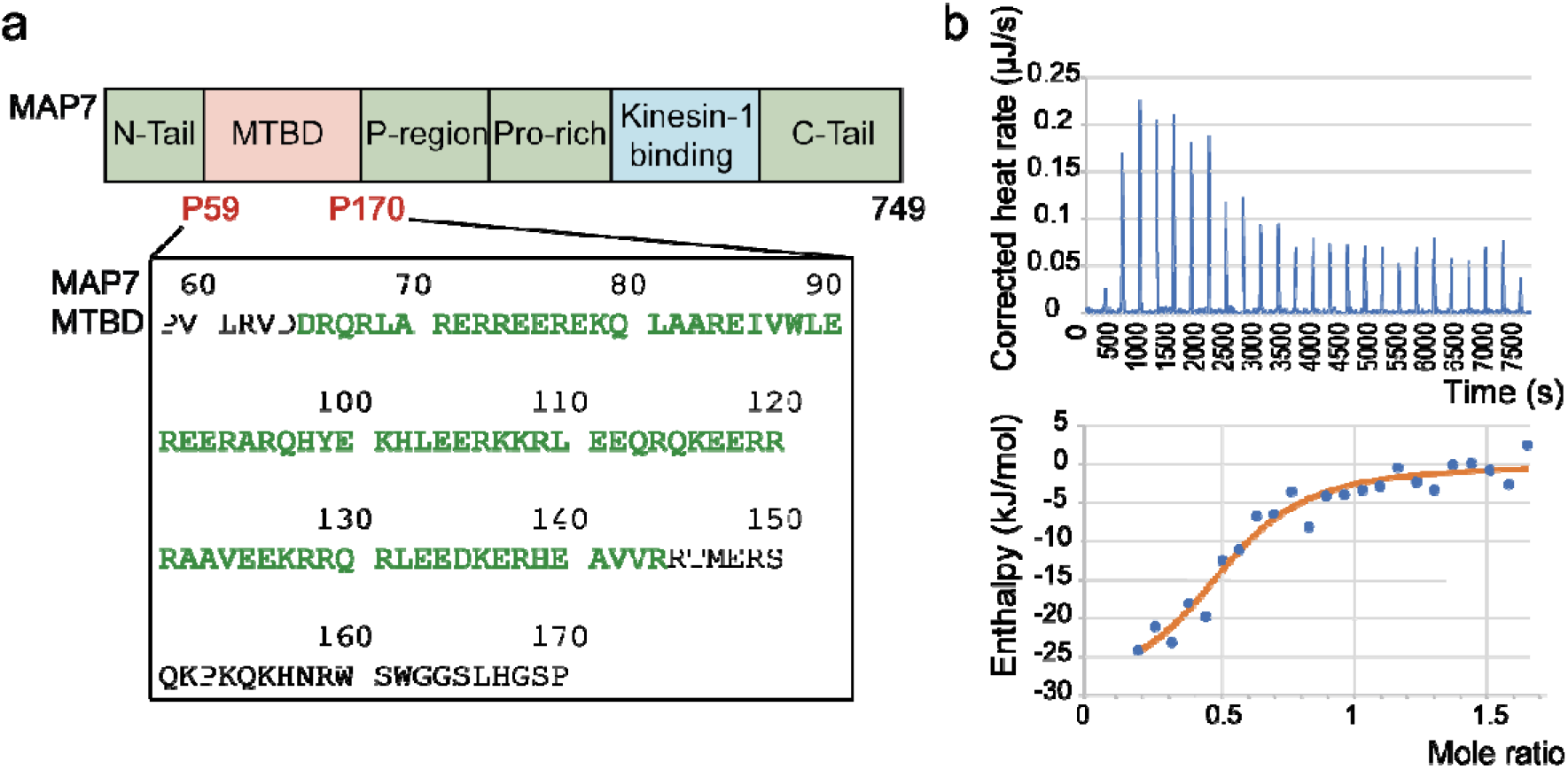
Binding of MAP7 MTBD to MT. **a** Top: Schematic representation of full-length MAP7 including the MTBD and the kinesin-1 binding domain. Bottom: Amino-acid sequence of the MAP7 MTBD, residues that are predicted to be α-helical are coloured in green ^17^. **b** Representative isothermal titration calorimetry (ITC) data of MAP7 MTBD to Taxol-stabilized MT. The top right picture displays the titration thermogram as heat release after each injection of MAP7 MTBD to the MT. In the lower right picture, the dependence of released heat in each injection is plotted against the ratio of MAP7 MTBD to MT.

In this study, we establish a comprehensive picture of the binding of MAP7 MTBD to MTs by utilizing a combination of nuclear magnetic resonance spectroscopy (NMR) and electron microscopy (EM), supported by fluorescence anisotropy and isothermal titration calorimetry experiments. Our results suggest an interaction with micromolar affinity between an extended α-helix and two tubulin dimers; this expands the previously determined MAP7-MT binding interface by around 25 residues, leaving only the N- and C- termini of the MTBD dynamic and explaining MAP7-promoted MT stabilization. We found that binding of MAP7 MTBD stabilizes the MT lattice via longitudinal interactions along the protofilaments. In addition, we employed solution-state NMR titration experiments using peptides comprising the tubulin CTTs to study the dynamic interactions between MAP7 MTBD at residue-specific level. We identified two important regions in MAP7 MTBD that interact with the CTTs and might be critical for recruitment of MAP7 on the MT lattice. Taken together, our experiments reveal how both strong as well as weak dynamic protein-protein interactions organize formation of the MAP7-MT complex.

## Results

### MAP7 MTBD binds with micromolar dissociation constant to MTs

We analyzed the binding of MAP7’s 112 residue long MTBD **(Fig. 1a)** to MT using solution-state NMR titrations of MTs to labelled MAP7 MTBD and isothermal titration calorimetry (ITC). Strong signal decrease in N-H-transverse relaxation-optimized spectroscopy (NH-TROSY) experiments at sub-stoichiometric amounts of added tubulin hints at a dynamic mode of binding with significant interconversion between free and MT-bound MAP7 MTBD **(Supplementary Fig. 1a and 1b)**. For the ITC experiments, MAP7 MTBD was titrated onto the MTs and a K_D_ of 0.94 µM ± 0.73 µM could be observed with a stoichiometry of 0.51 ± 0.09 (MAP7 MTBD to tubulin) **(Fig. 1b).** These values are similar to previous results which observed a K_D_ of 1.39 ± 0.5 µM using Total Internal Reflection Fluorescence microscopy (TIRF-M) of MAP7ΔC (aa 1–353) in (30 mM HEPES pH 7.4, 150 mM K-acetate, 2 mM Mg-acetate, 1 mM EGTA, and 10 % glycerol) ^35^. At low ionic strengths (30 mM HEPES, 5 mM MgSO_4_, 1 mM EGTA, pH 7.0) a ∼10-fold higher affinity has been reported (K_D_ 111 ± 12 nM) ^18^ indicating a significant contribution of electrostatic interactions in MTBD-MT binding.

### MAP7 MTBD binds along the MT protofilament and stabilizes the MT lattice

We first used EM to directly visualize the effect of MAP7 MTBD interaction with MTs during polymerization. 5 µM tubulin (below its critical concentration) was incubated in the presence of GTP with different concentrations of MAP7 MTBD and the resulting MTs were imaged using negative stain EM. We observed an increase in MT number at molar ratios of 0.25:1 and 0.5:1 of MAP7 MTBD:tubulin, whereas no MTs were observed when MAP7 was not included. This demonstrates that MAP7 MTBD promotes MT polymerization **(Supplementary Fig. 2)**. In samples with tubulin and MAP7 MTBD in stoichiometric amounts, bundling of MTs was observed, while at higher concentration ratios of 2:1 for MAP7 MTBD and tubulin, thick-walled short MTs containing an additional layer of protein, potentially composed of tubulin, were observed **(Supplementary Fig. 2).**

To further understand the MAP7 MTBD–MT interaction, and based on the distribution and number of MTs formed in these samples, we used the ratio of 1:1 (MAP7 MTBD:tubulin) to polymerize MTs for cryo-EM experiments. Using our established reconstruction pipeline ^36,37^ and treating the α/β-tubulin dimer as the asymmetric unit of the reconstruction, we obtained a symmetrized 3D reconstruction of MAP7 MTBD bound MT with an overall resolution of 3.7 Å which showed extra density corresponding to MAP7 MTBD along the protofilaments **(Supplementary Fig. 3a)**. However, the density corresponding to MAP7 MTBD, was only visible at inclusive map thresholds. To further improve the occupancy of MAP7 MTBD on the MT lattice, additional MAP7 MTBD was added to the stabilized MTs adsorbed on the grid before vitrification. The resulting symmetrized 3D reconstruction of MAP7 MTBD bound MT thus obtained had an overall resolution of 3.5 Å and revealed improved density corresponding to MAP7 MTBD at each protofilament crest, ∼ 20 Å from its centre **(Fig. 2a, Table 1, Supplementary Fig. 3b)**. To facilitate interpretation, we also used AlphaFold 2 multimer ^38^ to predict the structure of MAP7 MTBD bound to a tubulin dimer **(Fig. 2b, ribbon model)**. The predicted model illustrated that the helical part of the MAP7 MTBD extends beyond a tubulin dimer, and its precise mode of interaction is therefore obscured through averaging in our reconstruction. Information about the extended MTBD-MT interaction could not be deconvoluted despite efforts to apply modern cryo-EM processing strategies ^39,40^. This, together with the *a priori* probability of MAP7 MTBD interacting out of phase with tubulin dimers along and between protofilaments around the MTs, resulted in lower resolution of the MAP7 MTBD density (∼4 Å) compared to the rest of the structure, and an inability to visualize the MTBD boundaries **(Supplementary Fig. 3c)**. We nevertheless, fitted the model with the highest confidence score (pTm+ipTM=0.88) into the electron density map to gain insight into this interaction **(Fig. 2b)**. MAP7 MTBD forms a long helix that spans ∼13 nm and its position in the predicted model aligned well with the corresponding density in the map confirming its binding position and helical conformation **(Fig. 2b)**. The binding site and register of the MAP7 MTBD helix on the MTs also matches that obtained from 3D reconstruction of MAP7 MTBD bound to Taxol stabilized-MTs^18^, demonstrating that MAP7 interacts with MTs in the same way independently of how the MTs are stabilized **(Supplementary Fig. 3d).** Further, extension of density for both α- and β-tubulin’s H12 helix could be observed in the reconstruction and additional C-terminal residues Val437-Ser439 for α-tubulin and residues Ala428-Thr429 for β-tubulin were modelled into the density **(Table 2)**. The ordering of these usually disordered residues could be explained by an interaction between the tubulin CTTs and MAP7; however, the direct interaction is not visible in our reconstruction and is presumably less ordered **(Fig. 2c)**.

**Fig. 2:**
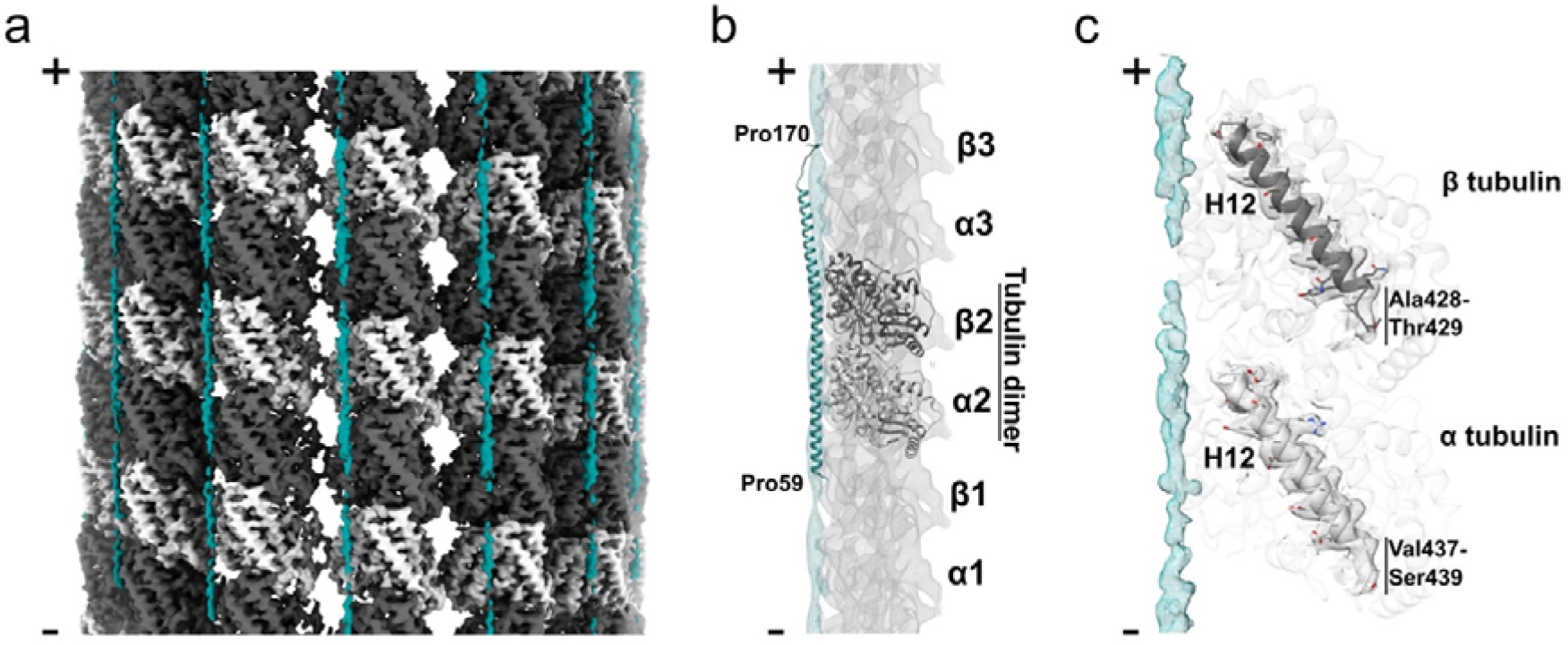
Cryo-EM reveals the binding site of MAP7 and its interactions along MT protofilaments. **a** Cryo-EM density (symmetrized reconstruction) of MAP7 MTBD bound MT. Density for the α-tubulin, β-tubulin and MAP7 MTBD is shown in light grey, dark grey and teal respectively. **b** Segment of symmetrized cryo-EM reconstruction corresponding to a protofilament, low-pass filtered to 10 Å shown in mesh representation with atomic model of MAP7 MTBD (teal) bound tubulin dimer (alpha tubulin: light grey; beta tubulin: dark grey) predicted by AlphaFold2 multimer fitted in. Position of tubulins along the protofilament and terminal residues of MAP7 MTBD have been indicated**. c** Density corresponding to H12 helix in both α- and β-tubulin as well as MAP7 MTBD helix shown in mesh representation with fitted atomic models for α- and β-tubulin depicted in cartoon representation. Residues that form the extension of the C-terminus from H12 helices are indicated in both α- and β-tubulin. Minus (-) and plus (+) ends of the MT are indicated in all panels.

**Table 1.** Data collection and image processing statistics.

|  |  |
| --- | --- |
| Microscope | Titan Krios D3771 |
| Voltage | 300 kV |
| Detector and mode | K3 detector and super-resolution counting mode |
| Pixel size (Å) | 1.067 |
| Defocus range (µm) | -0.6 to -2.4 |
| Software used | EPU |
| Total electron dose (electrons/ Å<sup>2</sup>) | 49.37 |
| Dose rate (electrons/pixel/s) | 16.53 |
| Exposure time (s) | 3.4 |
| Frames per movie | 50 |
| Number of micrographs | 2190 |
| Symmetry | Pseudo-helical |
| Initial number of particles | 83471 |
| Final number of particles | 15857 |
| Initial model used | EMD-7973 |
| Resolution (Å, FSC=0.143) | 3.5 |
| Local resolution range (Å) | 3.3 - 5.2 |
| Map sharpening factor ($\text{\AA}^2$) | -50 |

**Table 2.** Model refinement statistics.

|  |  |
| --- | --- |
| Initial model used | Sequence based model from ModelAngelo |
| Model composition |  |
| Non hydrogen atoms | 6338 |
| Protein residues | 857 |
| Ligands | 3 |
| RMSD of bond lengths ($\text{\AA}$) | 0.003 |
| RMSD of bond angles ($^\circ$) | 0.551 |
| MolProbity score | 1.81 |
| Clashscore | 9.06 |
| C-beta outliers (%) | 0 |
| Rotamer outliers (%) | 0.17 |
| Ramachandran plot |  |
| Favoured (%) | 95.29 |
| Allowed (%) | 4.71 |
| Outliers (%) | 0.00 |
| CC (mask) | 0.81 |
| CC (volume) | 0.79 |

### MAP7 MTBD shows an extended α-helix and dynamic terminal regions upon MT binding

To further probe the MAP7 MTBD–MT complex at atomic resolution, we conducted dipolar and J coupling (scalar)–based magic-angle spinning (MAS) solid-state NMR experiments, which provide complementary information about complex biomolecules that comprise both rigid as well as dynamic protein domains ^19,41^ **(Fig. 3a).** For our studies, we prepared samples in which [^13^C-^15^N] labelled MAP7 MTBD was mixed with Taxol-stabilized MTs at a 1 to 2 ratio of MAP7 MTBD to tubulin.

**Fig. 3:**
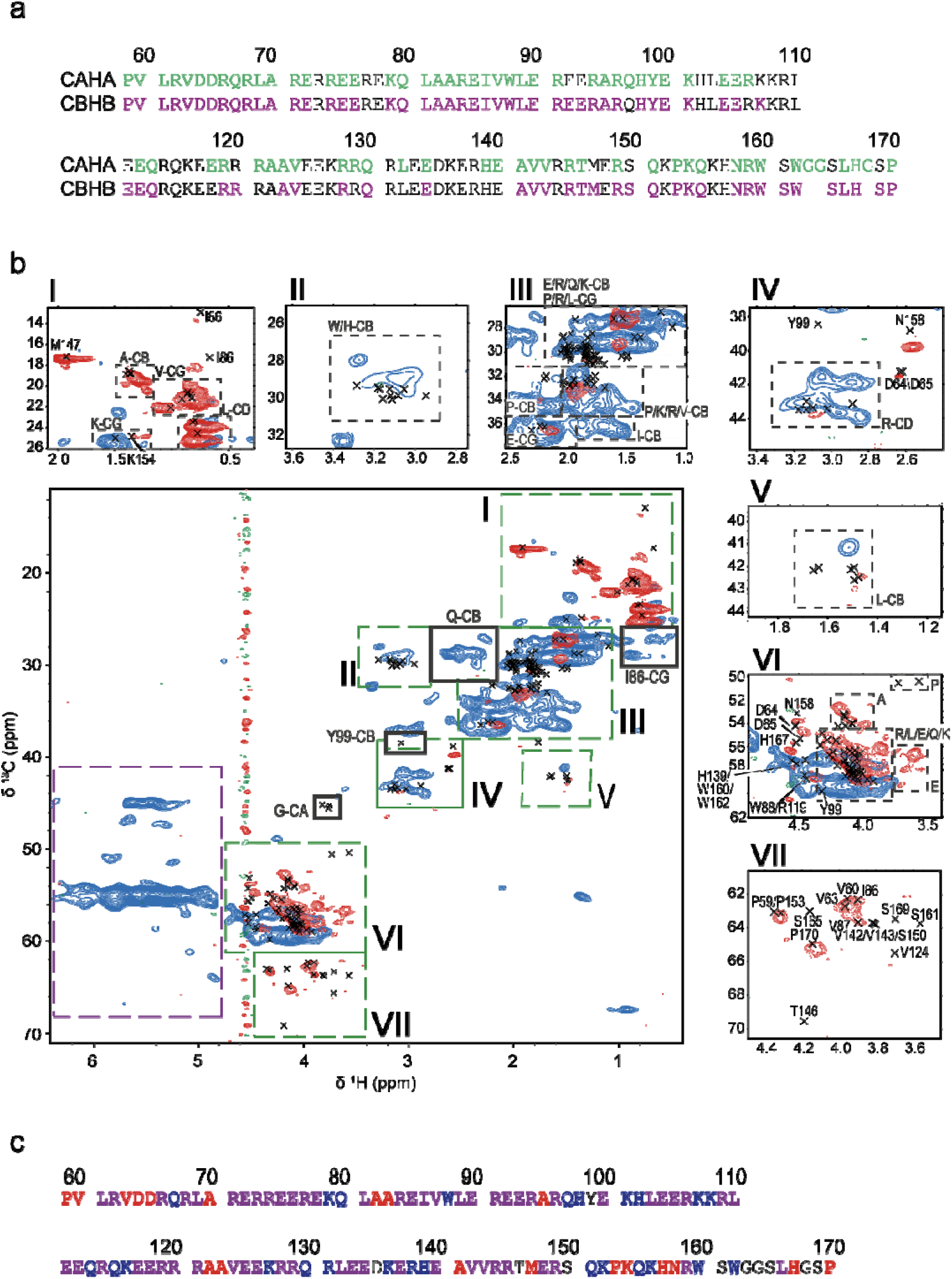
2D ssNMR of [^13^C-^15^N] MAP7 bound to Taxol-stabilized MT. **a** Solution-state assignments of MAP7 MTBD for CAHA (green) and CBHB (purple) resonances **b** Overlay of scalar (flexible) CH (red) and dipolar (rigid) CH (blue) on solution-state assignments of MAP7 MTBD (black crosses). Signal attributed to β-sheet chemical shifts is shown in the purple dashed box. Green roman numbers and boxes are used to indicate regions that are enlarged. **c** Summary of the observations from the 2D NMR spectra. In red are residues that overlay with the scalar CH, in blue those that overlay with the dipolar and in purple those overlaying with both.

The two-dimensional ^13^C-^1^H correlation experiments give a fingerprint of the rigid **(Fig. 3b, blue)** and flexible **(Fig. 3b, red)** components of the MAP7 MTBD bound to MTs. For reference, the solution-state assignments of MAP7 MTBD **(Supplementary Table 1** ^17^) are included as black crosses. A comparison of these data sets strongly suggests that the dipolar- and scalar-based ssNMR experiments probe different domains of MAP7. For further analysis, we made use of our previous solution NMR assignments which cover 97 % of CA and 93 % of CB resonances as well as 73 % of the HA and 69 % of the HB assignments **(Fig. 3a, Supplementary Table 1)**. Because of spectral overlap, missing assignments in side-chains and chemical shift perturbations, a direct transfer of the solution-state chemical shift assignments to the ssNMR spectra was hindered. Instead, we used in these cases average biological magnetic resonance data bank (BMRB) chemical-shift values **(Supplementary Table 1).**

Panels **(I)-(VII)** show that Pro, Val, Ala, Asp, Met and Asn residue types are largely dynamic, whereas Gln, Trp and Lys are mostly found to be rigid. Residues Lys154, His167 were additionally identified to be dynamic. Most other residue types revealed ssNMR correlations in both rigid and dynamic protein regimes **(Supplementary Methods).** In summary, analysis of our 2D ssNMR data sets suggests that flexible residues are more likely to be located at the N- or C-terminus of the protein, while rigid residues are mainly found in the α-helical region of MAP7 **(Fig. 3c)**.

Interestingly, additional correlations appeared in our dipolar ssNMR spectra in the proton region between 4.7 and 6 ppm (**Fig. 3b, purple dashed box**), which cannot be explained by our solution NMR data or a protein species that comprises α-helical or random-coil conformations. Instead, these signals exhibit β-strand character (*vide infra*). Notably, aggregation of MAPs with MT stabilizing function has been observed for MAP1b, MAP2, TPPP and Tau ^42–44^.

To validate these findings and to obtain further residue-specific information we conducted three-dimensional CCH solid-state NMR spectra **(Fig. 4, Supplementary Fig. 4a)** using both dipolar as well as scalar-based transfer steps.

**Fig. 4:**
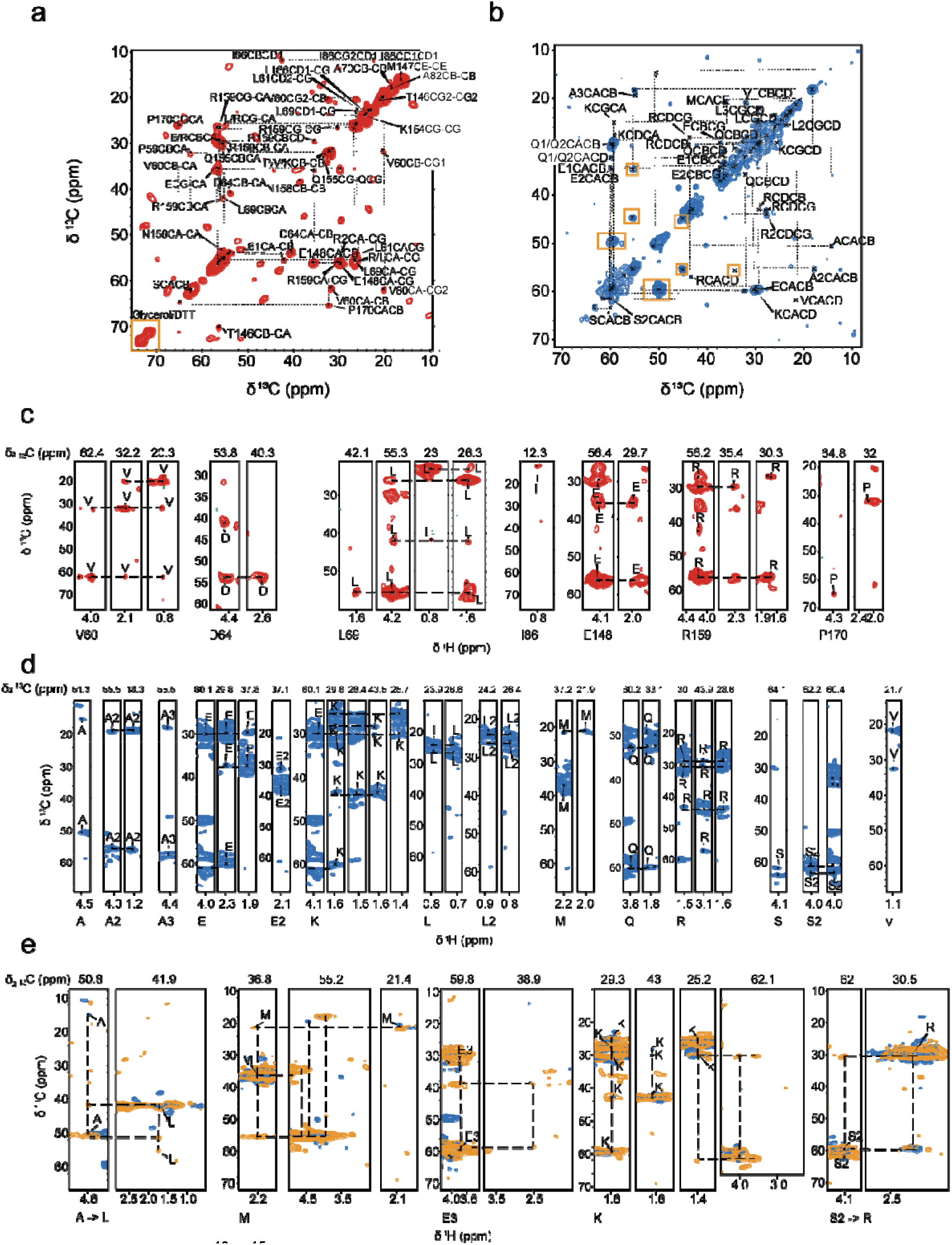
3D ssNMR of [^13^C-^15^N]-MAP7 bound to Taxol-stabilized MT. **a** CC-plane of 3D 11 ms DIPSI with assigned resonances. In the yellow box is signal produced by buffer components. **b** CC-plane of 3D 1.7 ms RFDR with assigned resonances. Yellow boxes indicate chemical shifts coming from the β-sheet signal. **c** Assigned strips of the scalar 3D CCH. **d** Assigned strips of the dipolar 3D CCH. **e** Assigned strips in the dipolar spectrum for 3.4 ms RFDR mixing time (yellow) with sequential connectivity compared to 1.7 ms RFDR (blue).

The 3D scalar spectrum (probing dynamic MAP7 residues in the complex) showed good agreement with the solution-state assignments. Here, we were able to assign several residues of MAP7 MTBD in the MT-bound state **(Fig. 4a and 4c).** For the Pro 59-Asp 64 stretch, Leu 69, Glu 148, Pro 153, Gln 155, Asn 158, Arg 159 and Pro 170 resonances for both backbone and sidechains could be identified **(Fig. 4a and 5a),** while for Ala 70, Ala 82, Ile 86, Thr 146, Met 147, Lys 154 and Leu 166 only certain side-chain resonances were found **(Supplementary Methods).** Intriguingly, all of these residues are located at the C- and N-terminus of MAP7 MTBD **(Fig. 5a),** in line with weak or no binding to MT for MAP7 residues Pro 59-Ala 70 and Glu 148-Pro 170. In addition, several resonances corresponding to residue types could be identified in the scalar CCH 3D, namely, one Glu, Ser, Gln, Leu and two additional Arg **(Fig. 4a).** Furthermore, we normalized the number of resonances of each residue type present in the aforementioned terminal domains to its overall abundance in the entire sequence. Taking the information gained from the experimental signal intensity together with the observed residues we find our scalar based experiments to be in reasonable agreement with the MAP7 regions Pro 59-Ala 70 and Glu 148-Pro 170 **(Fig. 5b, Supplementary Fig. 4b, Supplementary Methods).**

**Fig. 5:**
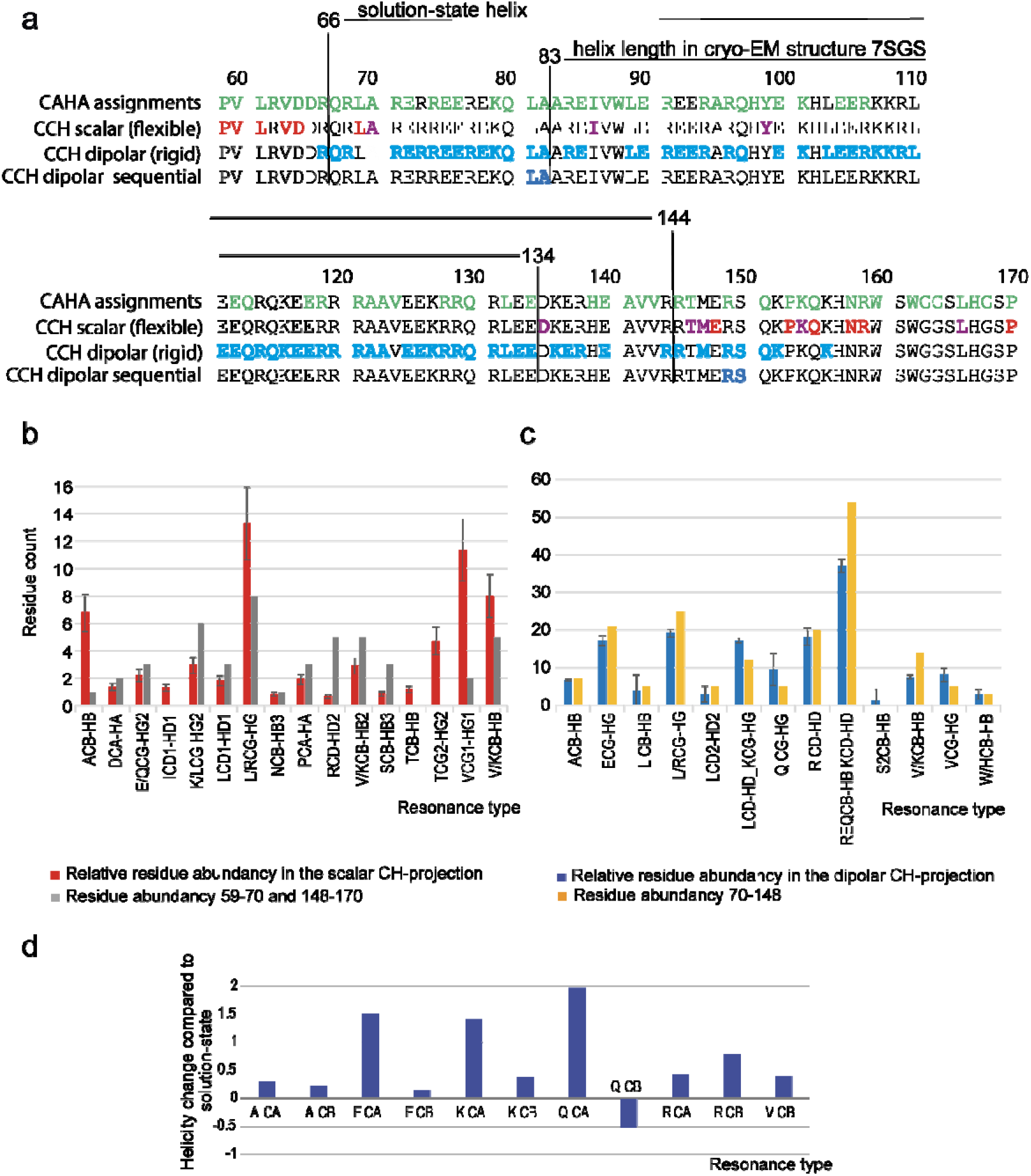
Analysis of 3D CCH ssNMR experiments. **a** Distribution of assigned residues indicated on the MAP7 MTBD sequence. Top: CAHA assignments in solution (green), 2nd row: assigned residues in the CCH scalar 3D (red for several resonances, magenta for only side-chain assignments), 3rd row: residue types found in dipolar CCH (blue), bottom row: residue pairs identified in sequential dipolar CCH. The length of the helix found in solution and in cryo-EM are indicated by black lines. The intensities for identified residue detected in scalar CCH **b** and the dipolar CCH **c** regions were integrated. The integrated intensities were divided by the signal average peaks attributed to one resonance only. The error bars have been propagated from the signal-to-noise ratio. **d** Comparison of helicity of solution-state assignment average for residue A70-E148 to average chemical shift helicity of the assigned same residue type in the dipolar CCH.

For the dipolar spectra **(Fig. 4b and d, Supplementary Fig. 4c)**, we were able to assign Ala, Glu, Lys, Leu, Met, Gln, Arg, Ser and Val residue types with distinct chemical-shift signatures **(Supplementary Table 2)**. These MAP7 MTBD residues are more abundant in the central α-helical region as seen in the prediction model fit into EM density and extend to approximately residue 150 **(Fig. 5a and 5c).** Indeed, these residue types exhibit α-helical shifts, underlining the α-helical secondary fold of the bound part of the MAP7 MTBD **(Fig. 5d).** It is interesting that Gln CBHB is an exception to the rule, as the chemical shift is more typical of random coil shifts than the average Gln CBHB chemical shift in solution. However, we identified only one Gln CBHB in the 3D CCH **(Supplementary Table 2),** which could be located in a region at the beginning or end of the helical part of MAP7 MTBD and was hence separated enough from the overlapping Gln resonances to be assigned (e.g.: Gln 80 or Gln 151).

We note that the aforementioned correlations are the ones that were unambiguously assignable. However, there are several spectral regions in which overlap precludes the assignment of individual resonances but still allows for a residue-type analysis. Similar to the scalar case, we calculated the relative abundance of these resonance-type specific regions by dividing the integral of the resonance region in the CH projection of the 3D CCH spectrum by the average of the integral of peaks corresponding to one assignment only **(Fig. 5c).** The residue abundance is in good agreement with the number of these residues in the region Ala 70-Glu 148 **(Fig. 5c, right and Supplementary Fig. 4b)**, corresponding to the domain that we previously found absent in the scalar spectrum **(Fig. 5a)**. This notion is confirmed by the presence of all expected Ala, Arg, Val, His, Gln and Trp side-chain resonances. Nonetheless, fewer Lys resonances were observed and none for Asp. The discrepancies may be due the fact that some of the residues exhibit a certain degree of motion which reduces dipolar transfer and cross-peak intensities in our dipolar ssNMR spectra.

In addition to the ssNMR data presented above, we also obtained sequential assignments from dipolar 3D CCH experiments recorded with a longer Radiofrequency driven dipolar recoupling (RFDR) mixing time (3.4 ms) **(Fig. 4e).** By comparing this 3D data set to results of the dipolar 3D experiment with shorter mixing time we could connect two residue pairs. Firstly, a sequential Ala-Leu contact would only be consistent with pairs 69-70 and 81-82. Because we identified Leu 69 in the scalar 3D, we tentatively assigned this correlation to Leu 81-Ala 82. Moreover, we observed sequential transfer from Ser to Arg which must stem from residues Arg 149 to Ser 150. Therefore, it seems likely that the rigid region extends up to approximately residue Ser 150, with possibly weaker binding starting from Thr 146 **(Fig. 4e and 5c).** The assigned resonances can be found in the supplementary table **(Supplementary Table 2).**

Similar to the analysis of the α-helical region, we also attempted a residue-specific analysis of the putative β-strand region. By verifying connections in the CAHA region of the dipolar spectrum corresponding to the aforementioned β-sheet resonances, we tentatively assigned residues that best agree with Ala 83, Arg 84, Glu 85, Ile 86 and Val 87 **(Supplementary Fig. 4b and Supplementary Methods)**. These residues would be in line with the aggregation propensity for MAP7 evaluated by AGGRESCAN, which identifies residues 84-89 to be aggregation-prone ^45^. Interestingly this region corresponds to the hinge, that was observed in the free MAP7 MTBD α-helix (residue 84-87) ^17^. The rest of the aggregate could not be observed in the NMR spectra, possibly due static conformational heterogeneity or intermediate exchange dynamics.

Taken together, our spectral analysis suggests a picture in which certain MAP7 residues are either flexible or rigid as depicted in **Fig. 5a**. We conclude that dynamics are prevalent in the N-terminus before residue 70 and in the C-terminus after residue 148.

### Two protein regions play a role in the MT carboxy-terminal tail interaction with MAP7 MTBD

Previous work hinted at the importance of the tubulin CTTs in binding of MAP7 to MTs as indicated by a reduction in the scalar signal of the CTTs ^19,33^. To examine the binding of MAP7 MTBD to the tubulin CTTs, we designed peptides comprising residues Ser 439-Tyr 451 (439-SVEGEGEEEGEEY-451) and Asp 427-Ala 444 (427-DATAEEEEDFGEEAEEEA-444) of HeLa S3 α- and β-tubulin, respectively. By titrating these to isotopically labelled MAP7 MTBD in solution **(Fig. 6a),** we were able to observe chemical-shift changes **(Fig. 6b).** Interestingly, we identified two main regions (82-AAREIVW-88) and (141-AVVRRT-146) that are affected by the presence of the CTTs peptides, while the latter region seems to be less modulated by the β-tubulin tail **(Fig. 6b).** Notably, we also detected other residues with lower sensitivity due to the presence of the CTT peptides, i.e., Asp 65, Glu 75, Arg 106, Ala 122 and Ser 150. We determined the binding affinity for the α-CTT with fluorescence anisotropy by attaching NHS-fluorescein to the peptide **(Fig. 6c).** The experiments resulted in a dissociation constant (K_D_) of 91.3 µM ±17.5. This binding affinity seems therefore to be 90 times weaker than the micromolecular interaction observed for binding of MAP7 MTBD to entire MTs **(Fig. 1b)**. The electrostatic character of the CTT–MAP7 interaction was validated by NMR measurements of MAP7 MTBD with 10-fold α-CTT at different salt concentrations. A reduction of chemical shifts due to salt increase could be observed, leading to a disappearance of any relevant chemical shift perturbations at 500 mM salt **(Fig. 6d).** The dynamic binding behaviour was also verified by previous total internal reflection fluorescence experiments showing a 4-fold reduction in the binding of the MAP7 MTBD to MTs upon removal of the CTTs via subtilisin cleavage ^18^.

**Fig. 6:**
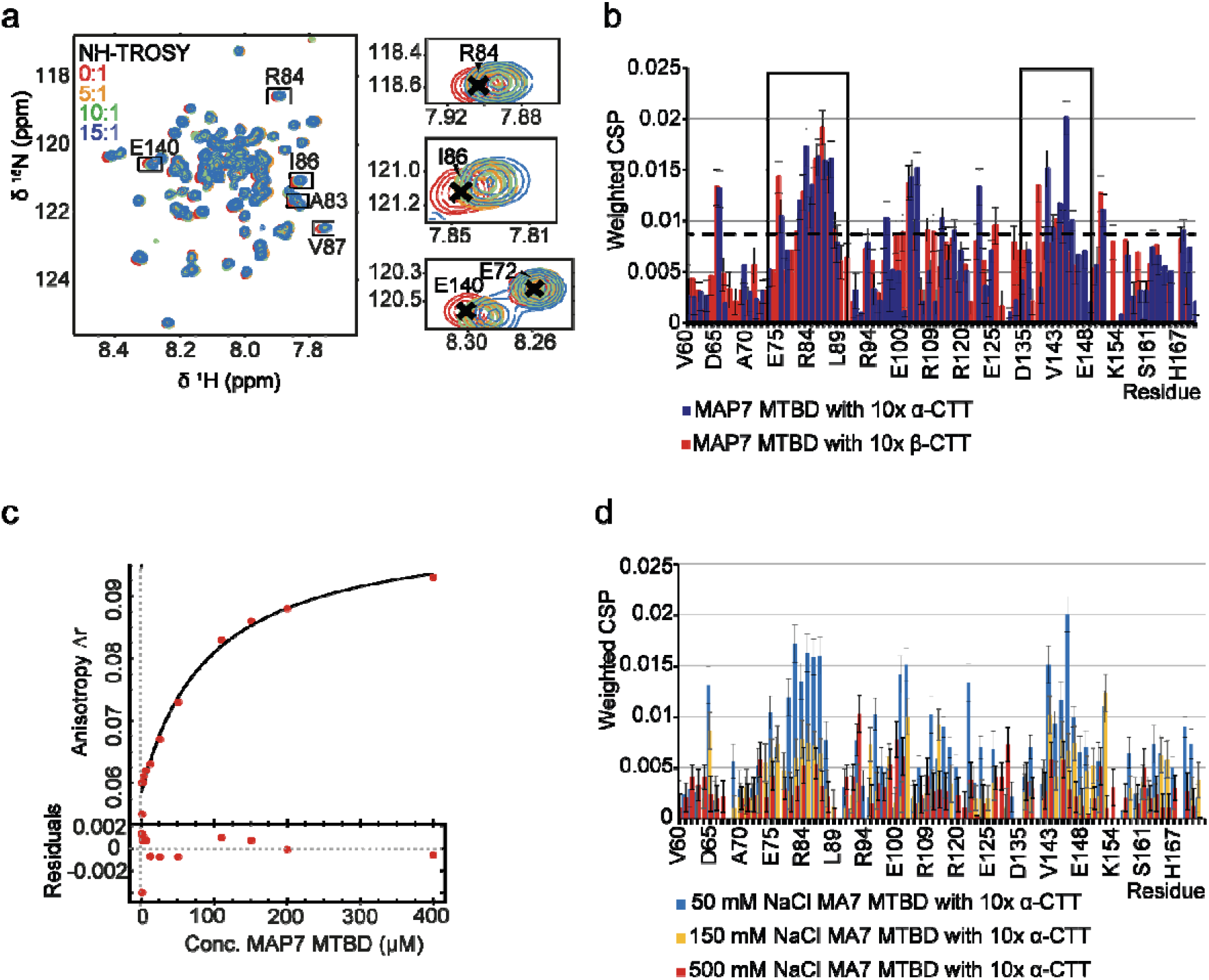
Titration of peptides comprising the MT carboxy-terminal peptides to [^13^C-^15^N]-MAP7 MTBD. **a** NH-TROSYs of the titration of increasing concentrations of α-CTT to MAP7 MTBD in solution-state NMR **b** Chemical shift perturbation (CSP) upon α- and β-CTT binding. Stretches with significant CSPs are marked with a box. The black dotted line indicates the mean plus one standard deviation**. c** Fluorescence anisotropy with fluorescein labelled α-CTT gives a KD of 91.3 µM ± 17.5 for the interaction with the MAP7 MTBD **d** CSP in labelled MAP7 MTBD upon addition of 10x α-CTT at different salt concentrations.

Our findings, in conjunction with the highly negative charge of the CTTs, suggest that the charged residues of the interacting stretches play a crucial role in an electrostatic binding event between MAP7 and MTs. Furthermore, the electrostatic character of the interaction is underlined by the observation that the FGE and EGE repeats of the CTTs are the binding residues on the MT side and therefore negatively charged residues might interact with the charged MAP7 residues ^19^.

## Discussion

MAP7 mediates recruitment of kinesin-1 to the MT lattice and plays a crucial role in the organization and transport of cellular cargo via MT-based motors, in organelle movement and spindle segregation ^46–48^. Here, we have studied the interaction between MAP7 MTBD and MTs by NMR and EM, allowing visualization of this interaction in the bound state. Despite the high repetitiveness of Arg, Glu, Lys residue types, we were able to acquire 2D and 3D solid-state NMR (ssNMR) spectra of labelled MAP7 MT-binding domain (MTBD) in complex with Taxol-stabilized MTs with remarkable spectral resolution. As confirmed by a combination of cryo-EM 3D reconstructions and AlphaFold2 prediction, MAP7 MTBD adopts an extended helical conformation when bound to MTs and its binding site on MTs when polymerized in presence of MAP7 MTBD is similar to that obtained when bound directly to Taxol-stabilized MTs. Our ssNMR experiments further identified the rigid and flexible parts of the protein, thereby establishing the helix boundaries and pinpointing specific interacting regions of the MAP7-MT interaction.

Because residues Arg 65-Arg 67 and Arg149-Ser150 are rigid, our data show that the MT binding interface of MAP7 extends beyond a single α/β-tubulin heterodimer **(Fig. 7 I. and IV.)**. The register is supported by the observation of more flexible residues in the clefts between the tubulin subunits **(Fig. 7 V. and VI.)** as well as by AlphaFold 2 multimer predictions, and is also consistent with a previous cryo-EM model ^18^. The resulting bridge between two tubulin dimers is predicted to support MT stabilization, as was shown when MTs were polymerized in sub-stoichiometric concentration ratios of 0.5:1 (MAP7 MTBD:tubulin) *in vitro* **(Supplementary Fig. 2)**, and is consistent with its observed stabilizing activity at axonal branches ^49^. The interaction of MAP7 with MTs is associated not only with MT remodelling but also kinesin-1-mediated transport in cells ^11^. As a result, studying this interaction will aid in understanding numerous cellular processes, which are linked to cancer-associated metastatic growth and neurodegenerative diseases ^12,35,50,51^.

**Fig. 7:**
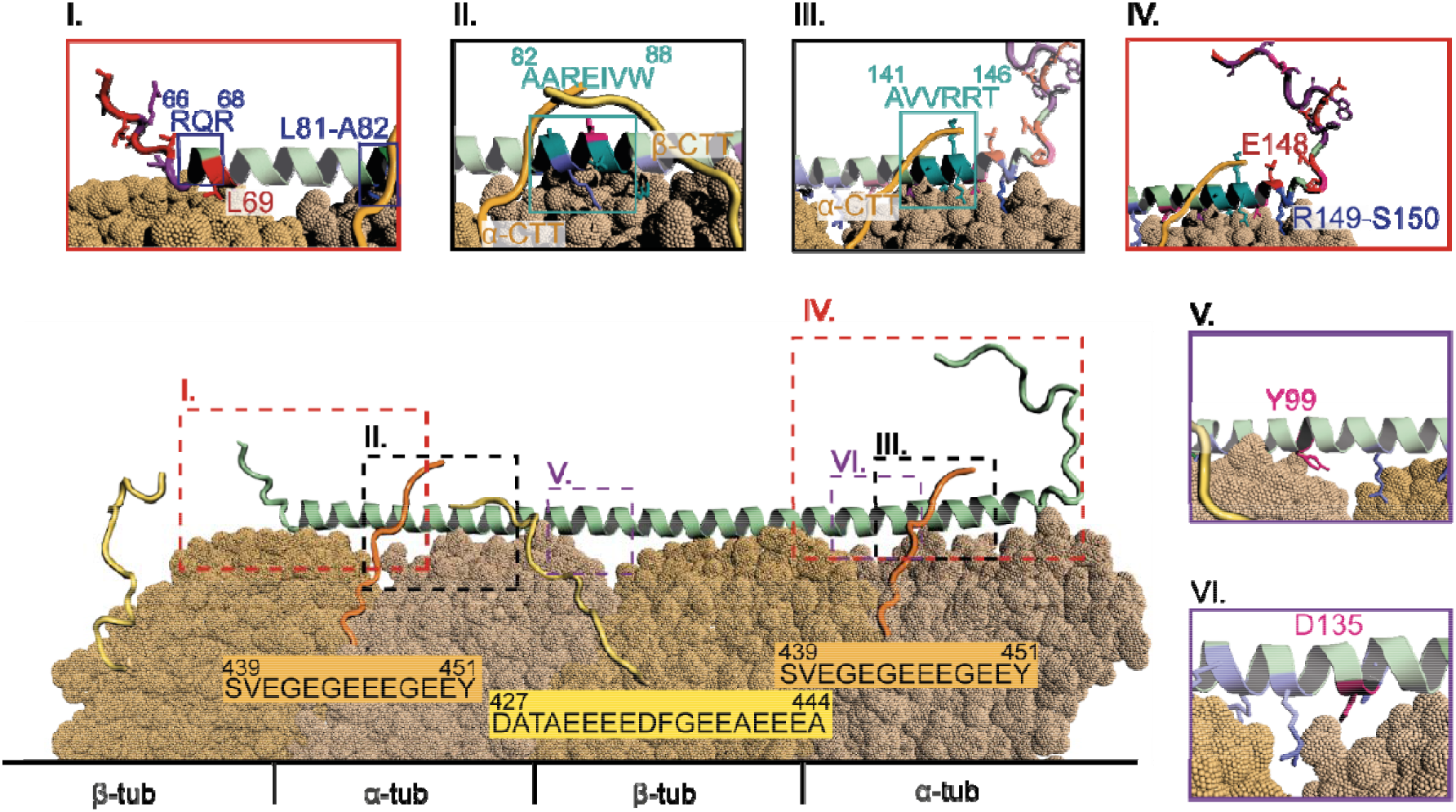
Model of MAP7 MTBD binding to MT based on the cryo-EM and NMR data. MAP7 MTBD binds MTs from around residue R66 to residue E148. On the N-terminal side backbone resonances with fast dynamics are only found prior to R66, with the exception of L69 and on the C-terminal side after E148 (red coloured, box **I.** and **IV.**). On the C-terminal side (panel **IV.**) a region with flexible and rigid residues is observed from T146 until S150. For T146 and M147 only side chain resonances are observed (pink). Box **II.** and **III.** show the interactions of MAP7 MTBD with the tubulin CTT. The residue stretches undergoing significant chemical shift perturbations are indicated by a teal square (box II.: A82-W88; box III.: A141-T146). I86 has flexible side-chains (pink, box II). In both cases, the CTTs are long enough to reach the identified MAP7 regions and also the residues of MAP7 on the neighbouring protofilament. Panels **V.** and **VI.** show residues with intermediate side-chain dynamics (pink) located between tubulins.

In addition to our earlier ssNMR results that revealed the binding site of MAP7 to the MT tails ^19^, solution-state NMR here provided insights into the dynamic interaction between tubulin’s carboxy-terminal tails (CTTs) and the MAP7 MTBD **(Fig. 7 II. And III.).** We identified two regions that displayed significant chemical shift changes. The first (residues 82 - 88) was found to interact with both the α-and β-CTTs, while the second region (residues 141-146) preferably interacted with the α-CTT. This interaction is further supported by the partial ordering of the tubulin C-terminal extensions of the H12 helix that were observed in the cryo-EM reconstructions. Together with previous studies that revealed the binding of MAP7 to the EGE and FGE motifs in MT CTTs ^19^, we now obtain an atomic-level understanding of the interaction between CTTs and MAP7. The long-range electrostatic interaction between the CTTs and the protein may help to attract MAPs from the cytoplasm to the MTs. Once binding has occurred, the relatively weak interaction between CTTs and MAP7 might contribute to stabilizing the MAP7-MT protofilament complex. This notion is also in line with observed weaker binding affinities of MAP7 to subtilisin-treated MTs ^18^.

Some ambiguities, such as whether the Arg residues in MAP7 involved in CTT binding are rigidified by the tail interaction or by the protofilament binding, cannot be clarified based on our ssNMR data. In addition, we cannot exclude the possibility that the CTTs may interact with the distal MAP7 bound to the neighbouring protofilament. Further studies with shorter MAP7 constructs might help to evaluate the importance of the N- and C-terminal interacting parts for MT stabilization. On the other hand, our approach could also be extended to study longer MAP constructs that are known to contain larger IDP regions ^52^, including MAP7 constructs containing the kinesin-1 binding domain. Finally, investigating post-translational modifications (PTMs) ^53^ of MT tails may provide insights into the importance of the “tubulin code” for binding affinities and binding rates leading to a more comprehensive understanding of these modifications and their implications in health and disease.

## Materials and Methods

### MAP7 MTBD expression and purification

The cDNA encoding the MTBD (residues 59-170) of MAP7 from *Homo sapiens* with an N-terminal His-Tag and Maltose binding protein (MBP) linked via a thrombin cleavage site (His-MBP-MAP7 MTBD), cloned into the pLICHIS vector was expressed in *Escherichia coli Rosetta 2 (DE3)* cells and purified with immobilized metal affinity chromatography, cation exchange chromatography, thrombin cleavage and ammonium sulphate precipitation ^17^.

### Negative stain sample preparation and EM

Porcine brain tubulin (Cytoskeleton, Cat No. T240) (5 µM) was incubated with MAP7 MTBD in BRB80 buffer (80 mM PIPES, 2 mM MgCl_2_, 1 mM EGTA, pH 6.8) and 2 mM GTP at indicated molar concentration ratios in a water bath maintained at 37°C for an hour. 3 µL of each sample was applied to glow discharged continuous carbon film coated, 400 mesh copper grids (Pacific Grid Tech) and blotted using a Whatman filter paper, Grade No. 1 after 30 seconds. Staining solution (2 % uranyl acetate in water) was then added to the grid, blotted off after 30 seconds and the grids were air dried. Images were recorded using Digital Micrograph™ software (Gatan) on a Tecnai T12 transmission electron microscope (Thermo Fisher Scientific) operating at 120 kV with a US4000 4K × 4K CCD camera (Gatan) at 3200x (overview) and 42000x (high) magnifications corresponding to pixel sizes of 13.5 nm and 2.5 Å respectively.

### Cryo-EM sample preparation

5 μM porcine brain tubulin (Cytoskeleton, Cat No. T240) and 5 μM MAP7 MTBD were mixed with 2 mM GTP (final concentration) in BRB80 buffer (80 mM PIPES pH 6.5, 2 mM MgCl_2_ and 1 mM EGTA) and incubated on ice for 5 minutes. The mixture was then incubated in a water bath maintained at 37 °C for an hour. C-Flat 2/2 Holey Carbon grids (Protochips) were treated in air for 40 seconds at 0.2-0.3 Torr using a Harrick Plasma Cleaner. 3 μl of the above sample was applied to and incubated on the surface-treated grid for 30 seconds at room temperature following which, excess sample was wicked off. For higher MAP7 occupancy and structure determination, 3.5 µL of 10 µM or 25 µM MAP7 MTBD was then immediately added to the grid. The grid was then mounted in a humidified Vitrobot Mark IV (Thermo Fisher Scientific) chamber pre-set to a temperature of 25 °C and humidity of 100 %. After incubation for 60 seconds, excess liquid was blotted out and the grid was plunge frozen in liquid ethane.

### Cryo-EM data collection

Data were collected using a Titan Krios D3771 microscope (Thermo Fisher Scientific) operating at an accelerating voltage of 300kV attached to a BioQuantum K3 direct electron detector (Gatan) and post-column GIF energy filter (Gatan) with a slit width of 20eV. The sample was imaged using the automated EPU software at a nominal magnification of 81,000x resulting in a pixel size of 1.067 Å and with defocus range of -0.6 to -2.4 μm. Movie frames were recorded in counting mode with a total dose of 49.37 electrons/ Å^2^ for 3.4 seconds fractionated over 50 frames.

### Cryo-EM image processing

Initial processing involved manual inspection of data for presence of contaminating ice and filament quality and poor-quality movies were discarded. All subsequent image processing steps were carried out using RELION v3.1 ^54,55^ and the customized Microtubule RELION-based Pipeline (MiRP) ^36,37^. Beam induced motion of particles in the movies were corrected using inbuilt function of MotionCor2 ^56^ in RELION. CTF estimation was performed on dose-weighted and motion corrected summed images using CTFFIND 4.1 ^57^ within RELION’s GUI. Particle picking was performed manually using RELION’s helical picker ^58^ and particles were extracted using a box size of 568 pixels and overlapping inter-box distance of 82 Å. The next steps of protofilament number segregation and Euler angle alignments were carried out using 2x binned particles and references with the help of MiRP v2 scripts^37^ integrated into RELION v3.1. MT segments with different protofilament numbers were segregated by supervised 3D classification using references for 11 to 16 protofilament containing MTs. 14 protofilament MTs formed the major fraction of the population and were re-extracted with a box size of 448 pixels for further processing. Rotational angle and X/Y coordinate fitting and shift assignments were carried out using 3D classification against a 15 Å low pass filtered reference (EMD-7973) with enhanced pixels for S9-S10 and H1-S2 loop regions in tubulin ^59^. Seam checking was carried out by supervised 3D classification using 28 rotated and shifted references. At this stage, aligned unbinned particles from 2 datasets (corresponding to 10 µM and 25 µM MAP7 externally applied) were combined. Finally, a symmetrized 3D reconstruction was obtained with a 3D auto-refine step using unbinned aligned particles and a 10 Å lowpass filtered reference **(Table 1)**. Per particle CTF refinement and Bayesian polishing were carried out in RELION and the final displayed reconstruction was sharpened using local resolution in RELION.

### Model generation

Protein sequences for porcine brain tubulin: porcine brain alpha-1A tubulin (UniPROT ID: P02550), beta-2B tubulin (UniPROT ID: P02554) and the symmetrized electron density map with a mask around the tubulin dimer in the centre of the protofilament opposite the seam were provided as inputs to locally installed ModelAngelo ^60^. The obtained model was further checked and refined manually by model building in Coot ^61^ followed by real-space refinement in Phenix ^62–64^ **(Table 2).** The quality of the electron density corresponding to MAP7 MTBD was insufficient to accurately build the MAP7 helix hence it was not modelled into the density. All images were made in Chimera^65,66^ or ChimeraX ^67,68^.

### AlphaFold2 multimer prediction

Protein sequences for porcine brain alpha-1A tubulin (UniPROT ID: P02550), beta-2B tubulin (UniPROT ID: P02554) and MAP7 MTBD (Residues 60-170 from UniPROT ID: Q14244) were used as inputs for locally installed AlphaFold2 multimer ^38^. Out of the 25 models obtained, the best model assessed by pTM+ipTM score was used for fitting and analyses. The model obtained was docked into the EM density using the ‘Fit in Map’ option of Chimera ^65,66^.

### Preparation of solid-state NMR samples

For the preparation of solid-state NMR samples lyophilized tubulin (Cytoskeleton, Inc.) was solubilised in BRB80 buffer (80 mM PIPES, 2 mM MgCl_2_, 1 mM EGTA, pH 6.8, 1 mM NaN_3_, 1 mM DTT, pH 6.8 supplemented with protease inhibitor (Sigma-Aldrich, cOmplete EDTA-free), to a final concentration of 2 mg/mL. Then 1 mM Guanosine-5’-triphosphate (GTP) was added and incubation took place for 15 minutes at 30 °C. In the following, 20 μM paclitaxel (Taxol, SIGMA) was used to stabilize the MT and incubation took place for another 15 minutes at 30 °C. The MT were spun down at 180,000 g (Beckman TLA-55 rotor) for 30 minutes at 30 °C and the pellet was resuspended in warm BRB80 buffer with 20 μM paclitaxel. Subsequently 0.55 mg/mL [^13^C-^15^N]-MAP7 MTBD was added. The interaction partners were incubated for 30 minutes at 30 °C. In the following step [^13^C-^15^N]-MAP7 MTBD in complex with MT was separated from the unbound, non-polymerised fraction by centrifugation at 180.000 g (Beckman TLA-55 rotor) for 30 minutes at 30 °C. Afterwards, the pellet was washed with BRB80 buffer containing protease inhibitor, without disturbing the pellet. A 1.3 mm rotor was packed with the pellet.

### Solid-state NMR

Solid-state NMR experiments on [^13^C-^15^N]-MAP7 MTBD with Taxol-stabilized MT were measured on a 700 MHz Bruker Avance III spectrometer with a 1.3 mm MAS rotor, at 55 kHz with a set temperature of 260 K, resulting in an effective sample temperature of approximately 299 K. All experiments were carried out with a ^1^H/X/Y triple-resonance MAS probe. Scalar-based 2D hCH were run and 3D hCCH correlation experiments were executed with 0 ms and 11 ms DIPSI ^69,70^ mixing time ^71^. Dipolar-based sequences were used with cross-polarization (CP) steps with an amplitude ramp of 80-100 % on ^1^H and 13 kHz PISSARO decoupling ^72^ during detection periods. CP transfer times were set to 700 µs forward CP and 150 µs back-CP. The dipolar 2D hCH and 3D hCCH experiments were recorded with 0 ms, 1.7 ms and 3.4 ms RFDR ^73^ mixing times. MISSISSIPPI was used for water suppression ^74^. All spectra were processed with the Bruker TopSpin 3.6.2 software. The data was zero-filled and an EM window function with a LB of 120 for the dipolar spectra and one of 30 for the scalar ones was applied. Linear prediction in the indirect ^13^C dimensions was utilized with acquisition times (ms) of direct-/indirect-dimension of 24/32/32 for the 3D dipolar, 30/20 for the 2D dipolar, 32/12 for the 2D scalar and 32/18/18 for the 3D scalar. Chemical shifts were referenced via the water resonance. The spectra were analysed using POKY from NMRFAM-Sparky ^75^. The residue abundance was estimated by signal integration of the peak position/region of interest and subsequent normalization to the integrated signal intensity average of well-resolved individual peaks that must represent a single resonance. For the scalar-based experiments, these resolved peaks refer to Ile 86 HG2CG2, Thr 146 CBHB, Asn 158 CBHB and Arg 159 HDCD resonances while to the dipolar data these correlations stem from Ser HBCB, Val CGHG, Leu CBHB and Ala CAHA. Resonance assignments for these reference peaks are highlighted in the **Supplementary Table 2**.

### Solution-state NMR

Solution-state NMR measurements for titration with MT were performed in NMR buffer (40 mM NaPi pH 6.5, 150 mM NaCl, 1 mM DTT) with 1 mM NaN_3_ and 1 mM DTT, in 10 % D_2_O. Two-dimensional (2D) ^1^H-^15^N TROSY experiments of starting concentration 30 µM [^13^C-^15^N]-labelled MAP7 MTBD, were recorded at 295 K on a 600 MHz Bruker Avance III spectrometer.

For the experiments with carboxy-terminal tails, peptides with the sequences SVEGEGEEEGEEY (α-CTT) and DATAEEEEDFGEEAEEEA (β-CTT) were purchased from Sigma-Aldrich and used without further purification. For the experiments the peptides were solubilised in NMR buffer and their pH adapted. For the NMR measurements increasing stoichiometric ratios of peptides to MAP7 MTBD were added to 80 µM [^13^C-^15^N]-MAP7 MTBD in NMR buffer. The 2D ^1^H-^15^N TROSY were recorded at 298 K on a 900 MHz Bruker Avance III spectrometer.

### Fluorescence Anisotropy

0.1 mg α-CTT was dissolved in 100 µL NMR buffer (40 mM NaPi pH 6.5, 150 mM NaCl, 1 mM DTT) and the pH adjusted to 8.3 by adding 30 µL of 0.5 M Na_2_HPO_4_.The concentration of the α-CTT was measured with a BCA assay (as previously mentioned). 40 mM NaPi, 150 mM NaCl, pH 8.3 was added to obtain a 500 µM α-CTT concentration. From here, the experiment was performed in the dark due to the light-sensitivity of NHS-fluorescein. 4 times molar ratio NHS-fluorescein to α-CTT was incubated for 2 hours at room temperature, inverted, whilst protected from light. The NHS-fluorescein-labelled α-CTTs was dialysed against NMR buffer (40 mM NaPi pH 6.5, 150 mM NaCl, 1 mM DTT) overnight at 4 °C in a 1 kDa cut-off dialysis tube, with a buffer exchange after 2 hours, to remove unbound NHS-fluorescein. By measuring the absorbance at 494 nm, combined with the extinction coefficient of 68/M*cm and the concentration of α-CTT from the BCA the concentration of labelled α-CTT was calculated. A dilution series of MAP7 MTBD resulting in a concentration range from 200 to 0 µM in a 384-well plate was prepared and NHS-fluorescein-labelled α-CTT to concentration of 1 µM was added to each well. We measured the fluorescence anisotropy at 494 nm with a plate reader and analysed the results with the DynaFit software (BioKin Ltd.) ^22^ .

### Isothermal titration calorimetry assay

Lyophilized tubulin (cytoskeleton) was solubilized in BRB80 buffer (80 mM PIPES, 2 mM MgCl_2_, 1 mM EGTA, pH 6.5, 1 mM NaN_3_, 1 mM DTT, pH 6.5 supplemented with protease inhibitor (Sigma-Aldrich, cOmplete EDTA-free)), to a final concentration of 2 mg/mL. Then 1 mM Guanosine-5’-triphosphate (GTP) was added and incubation took place for 15 minutes at 30 °C. In the following, 20 μM paclitaxel (Taxol, SIGMA) was used to stabilize the MT and incubation took place for another 15 minutes at 30 °C. The MT were spun down at 180.000 g (Beckman TLA-55 rotor) for 30 minutes at 30 °C and the pellet was resuspended in warm BRB80 buffer with 20 μM paclitaxel. The BRB80 buffer was then exchanged to warm NMR buffer with 1 mM β-mercaptoethanol (β-ME) instead of DTT and 1 mM GTP with a PD-10 desalting column.

Isothermal titration calorimetry was performed with the NanoITC (Waters LLC, New Castle, DE, United States) to determine the interaction between Taxol-stabilized MT and MAP7 MTBD in NMR buffer with 1 mM β-ME instead of DTT and supplemented with 1 mM GTP. Samples were degassed before performing the experiment. 150 µM MAP7 MTBD was titrated in 2 µL steps to 164 µL of 15 µM Taxol-stabilized MT. The protein was titrated at a rate of 2 μl/300 s with a stirring rate of 300 rpm. Experiments were performed at 37°C. Control experiments were performed with buffer titration. The dissociation constant (K_D_) value was calculated using the Nano Analyse Software (Waters LLC).

## Data availability

The cryo-EM map is deposited in the EMDB with accession number EMD-17469 and the corresponding atomic model for the tubulin dimer is available in the PDB with accession number 8P6R. All other data supporting the findings of this study are available in the article and the Supplementary Information.

## Acknowledgments

Funding: This work was supported by NWO (the Dutch Science Foundation) via a TOP-PUNT (grant number 718.015.001) grant to M. B., by uNMR-NL, the National Roadmap Large-Scale NMR Facility of the Netherlands (grant number 184.032.207) and by Medical Research Council, U.K. (MR/R00352/1) to C.A.M.). We acknowledge Diamond Light Source for access and support of the cryo-EM facilities at the UK’s national Electron Bio-imaging Centre (eBIC) funder proposal EM20287-56, funded by the Wellcome Trust, MRC and BBSRC. Cryo-EM data for the final EM reconstruction were collected at ISMB EM facility (Birkbeck College, University of London) with financial support from the Wellcome Trust (202679/Z/16/Z and 206166/Z/17/Z). We thank N. Lukoyanova and S. Chen for electron microscope support and D. Houldershaw for computational support at Birkbeck. A. A. thanks Dr. U. B. le Paige for fruitful discussions and ideas.

## Author contributions

A.A. carried out protein purification, MT assemblies, the solid-state NMR measurements with the help of SB. A.A. and M.B. wrote the paper with input from all authors. M. Ban conducted the EM experiments. M. B. and C.M. supervised the project.

