## Supporting information for "A structural and dynamic visualization of the interaction between the microtubule-associated protein 7 (MAP7) and microtubules"

**Supplementary Figures 1-4**

**Supplementary Tables 1-2**

**Supplementary Methods**

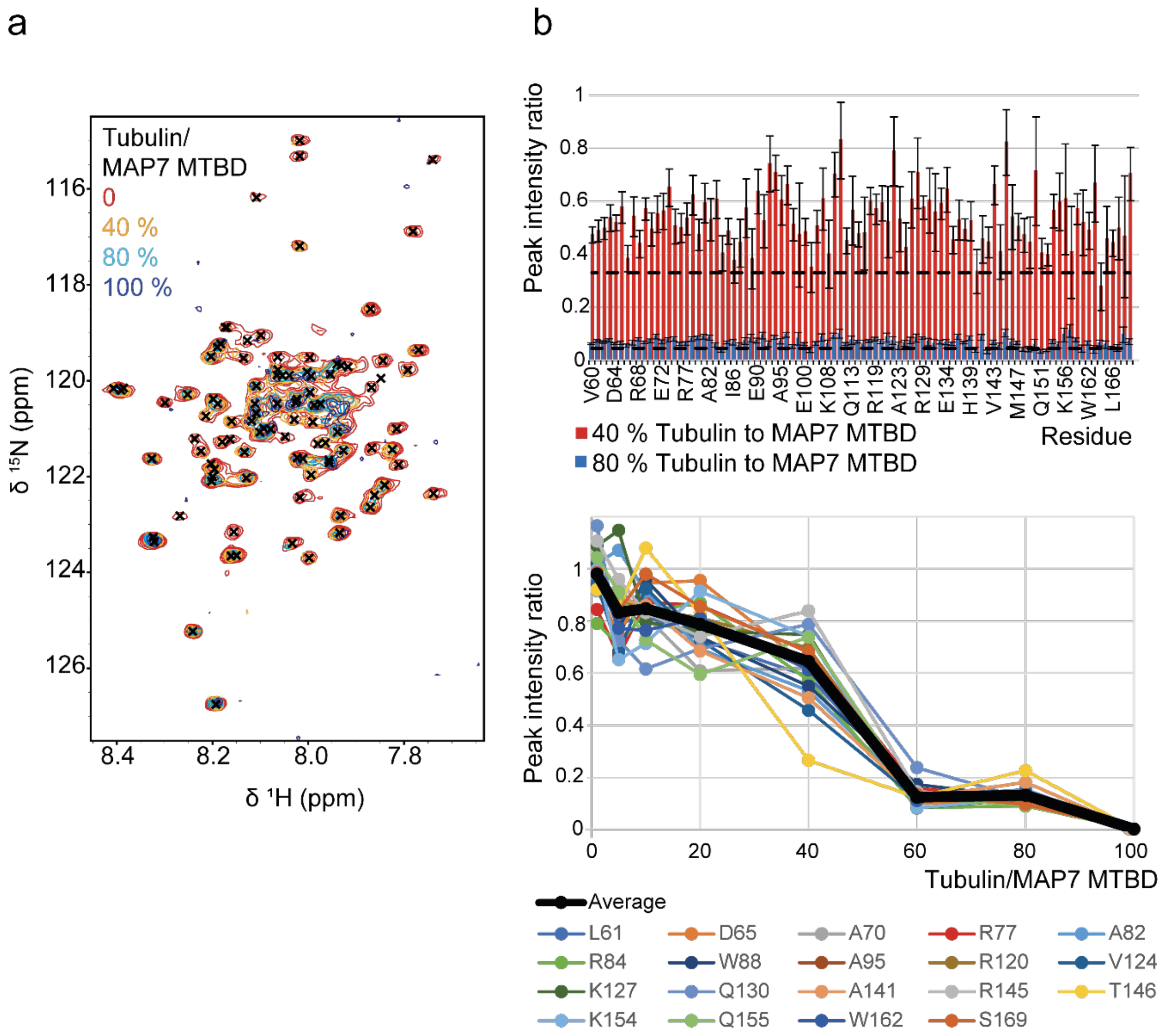


**Supplementary Fig. 1: Titration of Taxol-stabilized MT to [$^{13}C$-$^{15}N$]- MAP7 MTBD in solution a** Overlay of $^{1}H$-$^{15}N$ TROSY spectra of varying MT concentrations added to MAP7 MTBD. **b** Intensity ratio over whole MAP7 (top) upon adding 40 % or 80 % polymerized tubulin dimers to MAP7 MTBD. The error bars have been obtained from the signal-to-noise ratio. The black dotted lines are the 10 % trimmed average. Intensity ratio decreases over titration of selected residues (bottom).

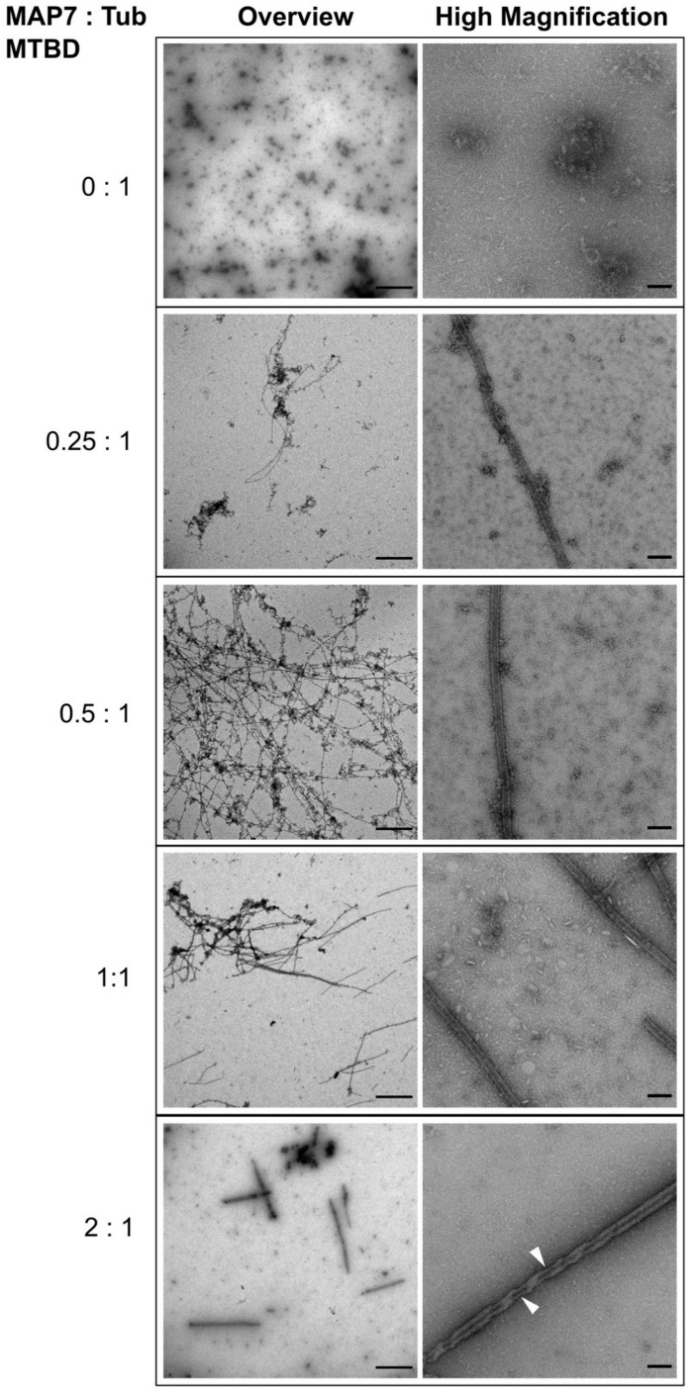


**Supplementary Fig. 2: Negative stain EM shows that MAP7 MTBD stabilizes dynamic MTs.** Left and right panels show micrographs at low (overview) and high magnifications respectively for MAP7 MTBD and tubulin incubated at molar concentration ratios of 0:1 (no MAP7 MTBD),0.25:1 , 0.5:1 , 1:1 and 2:1. Concentration of tubulin used is 5 μM. Scale bars represent 2 μm and 100 nm for the left and right panels respectively. White arrows in bottom panel (2:1 MAP7 MTBD: tubulin) show undecorated regions of MT lattice between the thicker surrounding protein (potentially tubulin) layer.

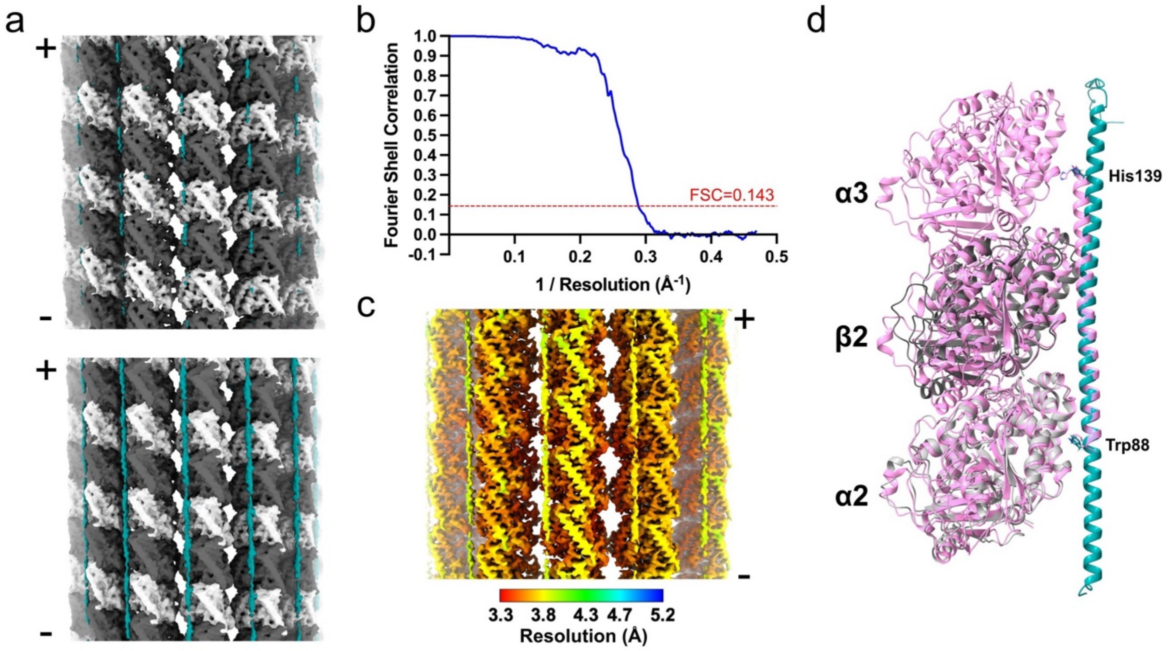


**Supplementary Fig. 3: Structure of MAP7 MTBD bound to MTs a** Cryo-EM density map (symmetrized reconstructions) of MAP7 MTBD bound MT (1:1 for MTBD and tubulin) when no MAP7 MTBD is added externally at stringent (top panel) and inclusive (bottom panel) map thresholds respectively. Density for the α-tubulin, β-tubulin and MAP7 MTBD is shown in light grey, dark grey and teal respectively. Minus and plus ends of the MT are indicated. **b** Fourier shell correlation plot (masked) for symmetrized reconstruction (Fig. 2a) of the MAP7 MTBD-MT complex. The resolution was estimated based on 0.143 criterion of half maps. **c** Cryo-EM reconstruction (Fig. 2a) coloured based on local resolution as determined by RELION with the colour scheme provided below. Minus and plus ends of the MT are indicated. **d** Cartoon representation of model predicted by AlphaFold2 multimer (same as Fig. 2b) superposed upon MAP7 MTBD:MT model (pink) previously determined by cryo-EM (PDB ID: 7SGS). Bulky residues corresponding to the ends of the 53-residue segment are shown as stick representation in both models and labelled.

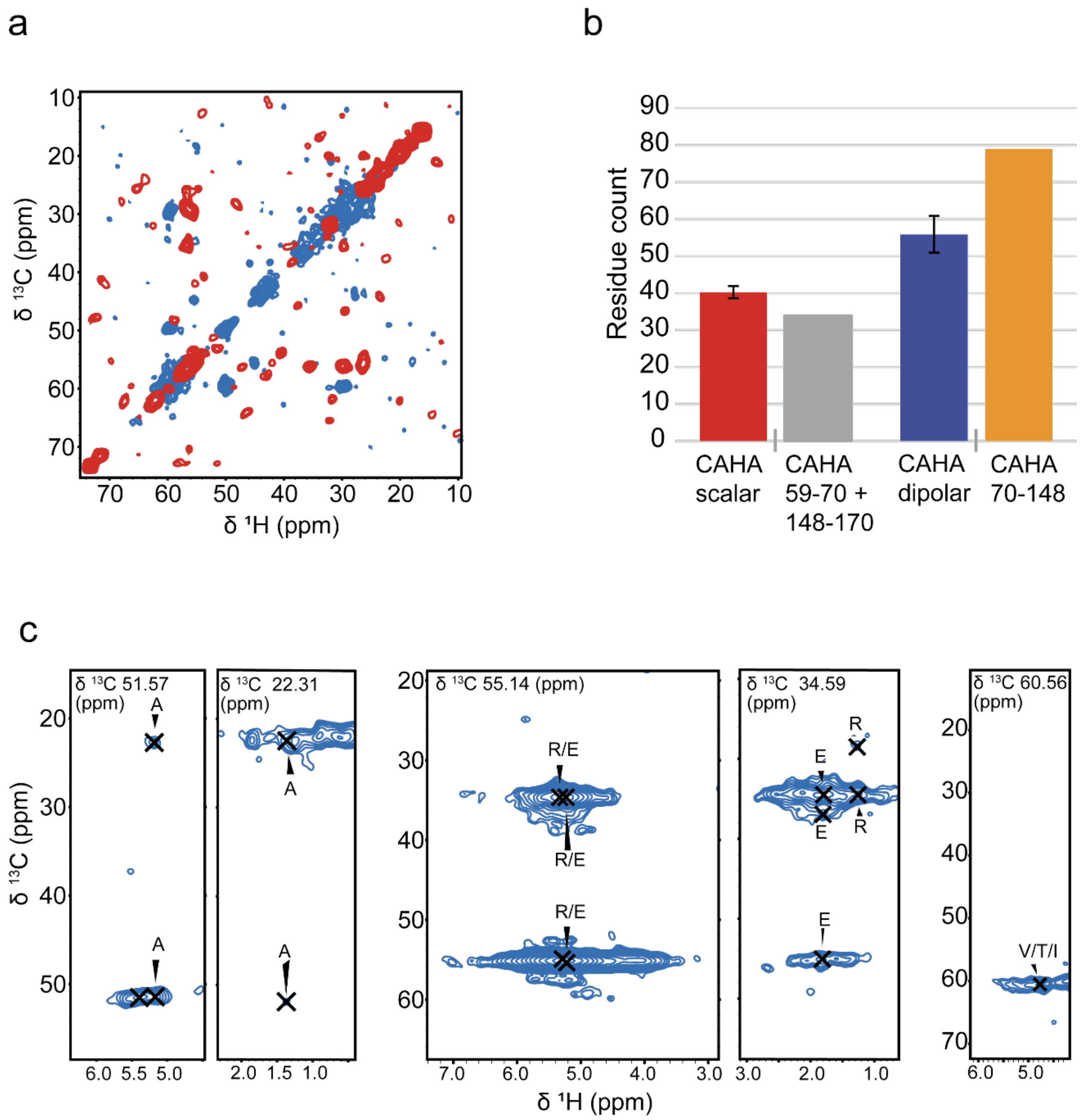


**Supplementary Fig. 4: 3D CCH experiments a** CC- planes of [$^{13}$C-$^{15}$N]-MAP7 MTBD bound to MT. Scalar and dipolar experiments are given in red and blue, respectively. **b** CAHA abundancy in MAP7 MTBD regions compared to peak integration of CAHA regions in the scalar and dipolar CH projections of the 3D CCH spectra. **c** Assigned strips of correlations in the dipolar 3D CCH that exhibit β-strand like chemical shifts.

### Supplementary Table 1: assigned chemical shifts from CH solution-state

| Assignment | $^{13}$C Shift | $^{1}$H Shift |
|---|---|---|
| P59CA-HA | 63.03 | 4.35 |
| P59CB-HB2 | 32.00 | 2.19 |
| P59CB-HB3 | 32.00 | 1.77 |
| V60CA-HA | 62.29 | 3.96 |
| V60CB-HB | 32.76 | 1.93 |
| V60CG1-HG1 | 21.14 | 0.83 |
| L61CA-HA | 54.88 | 4.32 |
| L61CB-HB3 | 42.59 | 1.49 |
| L61CD1-HD1 | 23.51 | 0.82 |
| R62CA-HA | 55.89 | 4.34 |
| R62CB-HB2 | 31.01 | 1.79 |
| R62CB-HB3 | 31.01 | 1.69 |
| R62CD-HD | 43.03 | 3.13 |
| V63CA-HA | 62.72 | 3.97 |
| V63CB-HB | 32.82 | 1.98 |
| V63CG1-HG1 | 20.64 | 0.86 |
| D64CA-HA | 54.23 | 4.53 |
| D64CB-HB2 | 41.24 | 2.63 |
| D64CB-HB3 | 41.24 | 2.61 |
| D65CA-HA | 55.32 | 4.49 |
| D65CB-HB2 | 41.23 | 2.62 |
| R66CA-HA | 58.36 | 4.01 |
| R66CB-HB2 | 30.08 | 1.83 |
| R66CD-HD | 43.51 | 3.13 |
| R66CG-HG | 27.21 | 1.53 |
| Q67CA-HA | 57.56 | 4.13 |
| Q67CB-HB2 | 28.71 | 2.04 |
| R68CA-HA | 58.26 | 4.03 |
| R68CB-HB2 | 32.71 | 1.98 |
| R68CB-HB3 | 32.71 | 1.78 |
| L69CA-HA | 56.56 | 4.09 |
| L69CB-HB2 | 42.04 | 1.64 |
| L69CB-HB3 | 42.04 | 1.49 |
| L69CD1-HD1 | 23.36 | 0.80 |
| A70CA-HA | 54.04 | 4.09 |
| A70CB-HB | 18.58 | 1.37 |
| R71CA-HA | 58.10 | 4.06 |
| R71CB-HB2 | 30.25 | 1.85 |
| R71CB-HB3 | 30.25 | 1.84 |
| R71CD-HD | 43.49 | 3.02 |
| E72CA-HA | 58.23 | 4.07 |
| E72CB-HB2 | 29.86 | 2.00 |
| R74CA-HA | 58.20 | 4.11 |
| R74CB-HB2 | 30.32 | 1.76 |
| R74CB-HB3 | 30.32 | 1.54 |
| R74CD-HD | 43.35 | 3.10 |
| E75CA-HA | 58.37 | 4.08 |
| E75CB-HB2 | 30.32 | 2.02 |
| E75CB-HB3 | 30.32 | 1.83 |
| E76CA-HA | 58.40 | 4.10 |
| E76CB-HB2 | 30.41 | 1.86 |
| E76CB-HB3 | 30.41 | 1.82 |
| K79CA-HA | 58.15 | 4.11 |
| K79CB-HB2 | 32.67 | 2.05 |
| K79CB-HB3 | 32.67 | 1.83 |
| K79CG-HG | 25.05 | 1.50 |
| Q80CA-HA | 57.16 | 4.14 |
| Q80CB-HB2 | 28.79 | 2.05 |
| Q80CB-HB3 | 28.79 | 1.83 |
| L81CA-HA | 56.40 | 4.11 |
| E100CA-HA | 58.28 | 4.07 |
| K101CA-HA | 58.37 | 4.09 |
| K101CB-HB2 | 31.00 | 1.84 |
| K101CB-HB3 | 31.00 | 1.78 |
| E104CA-HA | 58.81 | 4.08 |
| E104CB-HB2 | 29.57 | 1.86 |
| E104CB-HB3 | 29.57 | 1.83 |
| E105CA-HA | 58.32 | 4.08 |
| E105CB-HB2 | 29.77 | 2.05 |
| E105CB-HB3 | 29.77 | 2.01 |
| R106CA-HA | 58.99 | 3.92 |
| K107CB-HB2 | 32.51 | 1.84 |
| E111CB-HB2 | 29.47 | 1.99 |
| E111CB-HB3 | 29.47 | 1.85 |
| E112CA-HA | 59.03 | 4.04 |
| E112CB-HB3 | 29.46 | 1.84 |
| Q113CA-HA | 58.56 | 4.14 |
| Q113CB-HB2 | 28.49 | 1.98 |
| Q113CB-HB3 | 28.49 | 1.79 |
| E118CA-HA | 58.79 | 4.04 |
| R119CA-HA | 58.79 | 4.44 |
| R119CB-HB2 | 30.22 | 1.85 |
| R120CB-HB2 | 30.24 | 1.85 |
| R120CB-HB3 | 30.24 | 1.78 |
| R121CA-HA | 58.81 | 3.97 |
| A122CA-HA | 54.20 | 4.13 |
| A123CA-HA | 54.29 | 4.18 |
| A123CB-HB | 18.35 | 1.38 |
| V124CA-HA | 65.48 | 3.71 |
| V124CB-HB | 32.19 | 1.83 |
| V124CG2-HG2 | 22.13 | 1.01 |
| E125CG1-HG1 | 36.13 | 2.24 |
| R128CA-HA | 58.28 | 4.08 |
| R128CB-HB2 | 29.65 | 1.97 |
| R128CB-HB3 | 29.65 | 1.80 |
| R129CA-HA | 57.38 | 4.08 |
| Q130CA-HA | 57.60 | 4.21 |
| Q130CB-HB2 | 28.78 | 1.42 |
| Q130CB-HB3 | 28.78 | 1.37 |
| L132CA-HA | 56.73 | 4.07 |
| L132CD1-HD1 | 24.58 | 0.78 |
| E134CA-HA | 57.91 | 4.10 |
| E134CB-HB2 | 30.05 | 2.06 |
| E134CB-HB3 | 30.05 | 2.01 |
| H139CA-HA | 57.16 | 4.45 |
| H139CB-HB2 | 30.05 | 3.16 |
| H139CB-HB3 | 30.05 | 3.11 |
| E140CA-HA | 57.45 | 4.05 |
| A141CA-HA | 53.47 | 4.14 |
| A141CB-HB | 18.89 | 1.36 |
| V142CA-HA | 63.71 | 3.82 |
| V142CB-HB | 32.45 | 2.04 |
| V142CG1-HG1 | 20.79 | 0.87 |
| V142CG2-HG2 | 21.28 | 0.91 |
| V143CA-HA | 63.67 | 3.81 |
| V143CB-HB | 32.34 | 1.94 |
| R145CA-HA | 56.99 | 4.24 |
| R145CB-HB2 | 30.70 | 1.80 |
| R145CB-HB3 | 30.70 | 1.76 |
| R145CD-HD | 43.46 | 3.18 |

| | | |
|---|---|---|
| L81CB-HB2 | 42.15 | 1.66 |
| L81CB-HB3 | 42.15 | 1.50 |
| L81CG-HG | 27.25 | 1.60 |
| A82CA-HA | 53.28 | 4.15 |
| A82CB-HB | 18.82 | 1.37 |
| A83CA-HA | 53.35 | 4.15 |
| A83CB-HB | 18.83 | 1.36 |
| R84CA-HA | 57.05 | 4.05 |
| R84CB-HB2 | 30.63 | 1.98 |
| E85CA-HA | 57.38 | 4.15 |
| E85CB-HB2 | 30.11 | 1.97 |
| E85CG-HG | 36.50 | 2.31 |
| I86CA-HA | 62.34 | 3.90 |
| I86CB-HB | 38.36 | 1.77 |
| I86CD1-HD1 | 12.78 | 0.76 |
| I86CG1-HG1 | 27.92 | 1.10 |
| I86CG2-HG2 | 17.25 | 0.67 |
| V87CA-HA | 63.65 | 3.90 |
| V87CB-HB | 32.32 | 1.95 |
| V87CG1-HG1 | 20.88 | 0.83 |
| W88CA-HA | 58.45 | 4.46 |
| W88CB-HB2 | 29.42 | 3.29 |
| W88CB-HB3 | 29.42 | 3.18 |
| L89CA-HA | 56.38 | 4.07 |
| L89CB-HB3 | 42.48 | 1.48 |
| E90CA-HA | 58.04 | 4.03 |
| E90CB-HB2 | 29.78 | 1.99 |
| E90CB-HB3 | 29.78 | 1.85 |
| R91CA-HA | 58.15 | 4.02 |
| R91CB-HB2 | 30.41 | 1.98 |
| R91CB-HB3 | 30.41 | 1.80 |
| E92CB-HB2 | 29.70 | 1.95 |
| E93CB-HB3 | 29.68 | 1.83 |
| R94CA-HA | 56.73 | 4.26 |
| R94CB-HB2 | 30.25 | 1.78 |
| R94CB-HB3 | 30.25 | 1.48 |
| A95CA-HA | 54.01 | 4.09 |
| A95CB-HB | 18.85 | 1.40 |
| R96CA-HA | 58.18 | 4.02 |
| R96CB-HB2 | 30.24 | 2.00 |
| R96CB-HB3 | 30.24 | 1.85 |
| Q97CA-HA | 58.23 | 4.09 |
| H98CA-HA | 57.70 | 4.11 |
| H98CB-HB2 | 29.89 | 3.12 |
| Y99CA-HA | 59.84 | 4.32 |
| Y99CB-HB2 | 38.44 | 3.07 |

| | | |
|---|---|---|
| T146CB-HB | 69.57 | 4.18 |
| M147CB-HB2 | 32.83 | 1.99 |
| M147CE-HE | 16.90 | 2.00 |
| E148CG1-HG1 | 36.20 | 2.20 |
| R149CA-HA | 56.48 | 4.24 |
| R149CB-HB2 | 30.69 | 1.82 |
| R149CB-HB3 | 30.69 | 1.73 |
| S150CB-HB3 | 63.82 | 3.81 |
| Q151CA-HA | 55.46 | 4.28 |
| Q151CB-HB2 | 29.66 | 2.03 |
| Q151CB-HB3 | 29.66 | 1.89 |
| P153CA-HA | 63.14 | 4.32 |
| P153CB-HB2 | 32.18 | 2.21 |
| P153CB-HB3 | 32.18 | 1.79 |
| P153CD-HD | 50.41 | 3.57 |
| P153CG-HG | 27.32 | 1.94 |
| K154CA-HA | 56.49 | 4.15 |
| K154CB-HB | 32.98 | 1.68 |
| K154CG-HG | 24.87 | 1.35 |
| Q155CA-HA | 55.53 | 4.21 |
| Q155CB-HB2 | 29.78 | 1.91 |
| Q155CB-HB3 | 29.78 | 1.84 |
| N158CA-HA | 53.15 | 4.52 |
| N158CB-HB3 | 38.81 | 2.58 |
| R159CA-HA | 56.61 | 4.03 |
| R159CB-HB2 | 30.43 | 1.85 |
| R159CB-HB3 | 30.43 | 1.49 |
| R159CD-HD | 43.13 | 2.89 |
| R159CG-HG | 26.65 | 1.20 |
| W160CA-HA | 57.20 | 4.56 |
| W160CB-HB3 | 29.53 | 3.06 |
| S161CB-HB2 | 63.75 | 3.57 |
| W162CA-HA | 57.51 | 4.52 |
| W162CB-HB2 | 29.55 | 3.17 |
| W162CB-HB3 | 29.55 | 3.13 |
| G163CA-HA | 45.58 | 3.77 |
| G164CA-HA | 45.24 | 3.76 |
| S165CB-HB2 | 62.98 | 4.16 |
| H167CA-HA | 55.65 | 4.53 |
| H167CB-HB2 | 29.89 | 3.08 |
| H167CB-HB3 | 30.16 | 2.94 |
| G168CA-HA | 45.17 | 3.83 |
| S169CB-HB3 | 63.47 | 3.71 |
| P170CA-HA | 64.90 | 4.15 |
| P170CD-HD | 50.56 | 3.73 |
| P170CD-HD | 50.56 | 3.73 |

**Supplementary Table 2: assigned chemical shifts in CCH spectra.** Resonance assignments for the reference peaks of the residue abundance estimation are highlighted in bold.

| Dipolar CCH | | |
|---|---|---|
| Residue | Atom | Shift |
| A | CA | 50.25 |
| A | CB | 14.96 |
| A | HA | 4.57 |
| A2 | CA | 54.80 |
| A2 | CB | 18.23 |
| A2 | HA | 4.05 |
| A2 | HB | 1.33 |
| **A3** | **CA** | **54.91** |
| A3 | CB | 19.37 |
| **A3** | **HA** | **4.43** |
| E | CA | 59.72 |
| E | CB | 29.59 |
| E | CG | 36.53 |
| E | HB2 | 2.28 |
| E | HA | 3.98 |
| E | HG2 | 1.93 |
| E2 | CB | 31.06 |
| E2 | CG | 36.70 |
| E2 | HG2 | 2.10 |
| E3 | CB | 30.32 |
| E3 | CG | 34.26 |
| E3 | HG2 | 1.25 |
| E3 | HA | 3.78 |
| K | CA | 59.66 |
| K | CB | 31.69 |
| K | CD | 29.44 |
| K | CE | 43.06 |
| K | CG | 25.21 |
| K | HD2 | 1.57 |
| K | HA | 3.78 |
| K | HG2 | 1.86 |
| K | HB2 | 1.57 |
| K | HE2 | 1.70 |
| L | CD2 | 22.42 |
| L | CG1 | 28.56 |
| L | CD1 | 25.92 |
| L | HD1 | 0.93 |
| L | HD2 | 0.73 |
| L | HG | 0.80 |
| L2 | CB | 42.08 |
| L2 | HB | 1.57 |
| L3 | CD | 25.49 |
| L3 | HD1 | 0.79 |
| L3 | HD2 | 0.82 |
| **L4** | **CB** | **42.24** |
| **L4** | **HB** | **1.4** |
| M | CB | 36.53 |
| M | CE | 20.61 |
| M | HB2 | 2.20 |

| Scalar CCH | | |
|---|---|---|
| Residue | Atom | Shift |
| E1 | CA | 56.3 |
| E1 | CG | 35.7 |
| E1 | CB | 29.62 |
| E1 | HA | 4.1 |
| E1 | HB2 | 2.16 |
| E1 | HG3 | 1.96 |
| E/R1 | CA | 55.75 |
| E/R1 | CB | 30.23 |
| E/R1 | HA | 4.23 |
| L1 | CD2 | 23.17 |
| L1 | HD2 | 0.83 |
| P/V/K1 | CB | 32.67 |
| P/V/K1 | HB3 | 1.74 |
| Q1 | CA | 55.22 |
| Q1 | HA | 3.9 |
| R/L1 | CA | 55.93 |
| R/L1 | CD | 42.12 |
| R/L1 | CG | 26.5 |
| R/L1 | HG3 | 1.59 |
| S1 | CB | 62.28 |
| S1 | HB3 | 3.51 |
| L1 | CD2 | 23.24 |
| L1 | HD2 | 0.79 |
| R1 | CA | 56.2 |
| R1 | CB | 30.41 |
| R1 | CG | 26.49 |
| R1 | HG2 | 1.78 |
| P59 | CA | 62.28 |
| P59 | CB | 31.6 |
| P59 | HA | 4.33 |
| V60 | CA | 62.07 |
| V60 | CB | 31.73 |
| V60 | CG1 | 20.41 |
| V60 | CG2 | 20.26 |
| V60 | HA | 3.95 |
| V60 | HB | 1.97 |
| V60 | HG1 | 0.85 |
| L61 | CA | 54.35 |
| L61 | CB | 42.13 |
| L61 | CD2 | 22.84 |
| L61 | CG | 26.06 |
| L61 | HB | 1.61 |
| L61 | HG | 1.51 |
| D64 | CA | 53.98 |
| D64 | CB | 40.42 |
| D64 | HA | 4.47 |
| D64 | HB | 2.61 |
| L69 | CA | 55.34 |
| L69 | CB | 42.17 |

| | | |
|---|---|---|
| **M** | HE | 1.92 |
| **Q** | CA | 59.84 |
| **Q** | CB | 29.21 |
| **Q** | CG | 32.22 |
| **Q** | HB2 | 1.88 |
| **Q** | HA | 4.07 |
| **Q** | HG2 | 1.69 |
| **R** | CA | 57.01 |
| **R** | CB | 29.62 |
| **R** | CD | 42.98 |
| **R** | CG | 27.91 |
| **R** | HD2 | 3.11 |
| **R** | HB2 | 1.50 |
| **R** | HG2 | 1.58 |
| **R2** | CG | 28.20 |
| **R2** | CA | 59.45 |
| **R2** | HD2 | 3.11 |
| **R2** | HB2 | 1.65 |
| **R3** | HG2 | 1.58 |
| **S** | CA | 59.93 |
| **S** | **CB** | **61.97** |
| **S** | **HB2** | **4.18** |
| **S2** | CA | 59.84 |
| **S2** | CB | 62.21 |
| **S2** | HB2 | 4.01 |
| **V** | CB | 31.94 |
| **V** | CG | 21.47 |
| **V** | HG1 | 1.16 |
| **V2** | **CG** | **22.78** |
| **V2** | **HG1** | **0.99** |

| | | |
|---|---|---|
| **L69** | CD1 | 26.36 |
| **L69** | CD2 | 23.04 |
| **L69** | CG | 26.42 |
| **L69** | HA | 4.17 |
| **L69** | HD1 | 1.57 |
| **L69** | HD2 | 0.77 |
| **L69** | HG | 1.56 |
| **I86** | **CD1** | **11.81** |
| **I86** | CD2 | 22.5 |
| **I86** | CG2 | 12.15 |
| **I86** | **HD1** | **0.75** |
| **I86** | HG2 | 0.81 |
| **T146** | CA | 62.47 |
| **T146** | **CB** | **70.36** |
| **T146** | CG2 | 20.96 |
| **T146** | HA | 4.12 |
| **T146** | **HB** | **4.09** |
| **T146** | HG2 | 1.1 |
| **M147** | CA | 60.15 |
| **M147** | CE | 16.42 |
| **M147** | HE | 1.92 |
| **E148** | CA | 56.42 |
| **E148** | CB | 29.56 |
| **E148** | CG | 35.69 |
| **E148** | HA | 4.12 |
| **E148** | HG3 | 1.95 |
| **K154** | CD | 28.39 |
| **K154** | CG | 24.15 |
| **K154** | HD3 | 1.58 |
| **K154** | HG3 | 1.34 |
| **Q155** | CA | 55.27 |
| **Q155** | CB | 30.38 |
| **Q155** | CG | 33.32 |
| **Q155** | HA | 3.89 |
| **Q155** | HG3 | 2.2 |
| **N158** | CA | 53.54 |
| **N158** | **CB** | **38.5** |
| **N158** | HA | 4.57 |
| **N158** | **HB** | **2.59** |
| **R159** | CA | 56.11 |
| **R159** | CB | 30.37 |
| **R159** | **CD** | **42.69** |
| **R159** | CG | 26.87 |
| **R159** | HA | 4.16 |
| **R159** | HB2 | 2.22 |
| **R159** | **HD2** | **3.1** |
| **R159** | HG2 | 1.71 |
| **L166** | CB | 41.01 |
| **L166** | CD2 | 23.85 |
| **L166** | HD2 | 0.82 |
| **P170** | CA | 64.86 |
| **P170** | CB | 31.96 |
| **P170** | HA | 4.18 |
| **P170** | HB2 | 2.13 |

### Supplementary Methods

#### Spectral analysis of 2D and 3D ssNMR data sets

The analysis of scalar and dipolar 2D $^{13}C$-$^{1}H$ correlation experiments in **Fig. 3b** was performed on the basis of previous solution NMR assignments and average BMRB chemical-shift values. For example, in panel **(I),** we find good agreement between peak positions for the Ala, Val and Leu side-chain resonances seen in the complex using J-based ssNMR (red) and solution-state results (crosses). In the same vein, correlations corresponding to Met 147 CE-HE, Lys 154 CG-HG and two peaks corresponding to Ile 86 QG1-HG1 and Ile 86 CD1-QD1 (where Q stands for one peak for multiple protons) are visible. On the other hand, most Lysine CG-QG side-chain correlations seem to appear in a spectral region only detected in the dipolar spectrum. Likewise, His and Trp CH-HB resonances seem to only appear in panel (**II**) in the dipolar-based ssNMR data. Region **(III)** shows side-chain resonances of Glu, Arg, Gln, Lys, Val, Leu, and Pro. In this region, both dipolar and scalar signals can be seen. Interestingly, Ile 86 CB-HB correlations seen in solution appears rigid (Blue, indicate in panel **III**), while as mentioned above for panel (I), Ile CD-QD and CG-QG might be flexible as there is only one Ile residue in the sequence. Region **(IV)** contains Arg CD-QD, Asp CB-HB, Tyr 99 CB-HB and Asn 158 CB-HB resonances. With some exceptions, MAP7 MTBD seems to have rigid Arg CD-QDs, suggesting stabilization of arginine sidechains after complex formation. On the other hand, the solution-state NMR CB-HB resonances associated with Asp 64, Asp 65 and Asn 158 are in good agreement with the scalar signals in the complex. The Leu CB-HB region that is depicted in panel **(V)** gives scalar, as well as dipolar signals. Panel **(VI)** shows the crowded CA-HA region. There are no resonances found for Pro CD-QD. Flexible CA-HAs include Ala CA-HA, several Glu CA-Has and CA-HA of Asp 64, Asp 65, Asn 158 and His 167. For the crowded Arg, Glu, Leu, Gln, and Lys region both dipolar and scalar signals appear. Lastly, panel **(VII)** depicts well-dispersed Pro, Ser, Val, Ile CA-HA resonances that only appear in the mobile ssNMR experiments and correlate well with the solution assignments of free MAP7. Particularly, Pro 59, Val 60, Ile 86, Val 87, Pro 153, and Pro 170 overlap. In addition, it is intriguing to note that the Gln CG-HG region is mainly visible in the dipolar spectrum.

#### 3D ssNMR data analysis

For the analysis of the 3D scalar spectrum (probing dynamic MAP7 residues in the complex) we observed good agreement with the solution-state assignments. Therefore, we were able to assign several residues of MAP7 MTBD in the MT-bound state **(Fig. 4a and 4c).** For Pro 59, Val 60, Leu 61, Val 63, Asp64, Leu 69, Glu 148, Pro 153, Gln 155, Asn 158, Arg 159 and Pro 170, resonances for both backbone and sidechains could be identified **(Fig. 4a and 5a)** while for Ala 70 and Ala 82 only CB resonances were found. In addition, we could tentatively assign Ile 86 CG and CD, Thr 146 CA-CB, CG, Met 147 CA-CE, CE; for Lys 154 CD-CG and Leu 166 CD-CG. Intriguingly, all of these residues are located at the C- and N-terminus of MAP7 MTBD **(Fig. 5a),** in line with weak or no binding to MT for MAP7 residues Pro 59-Ala 70 and Glu 148-Pro 170. In addition, several resonances corresponding to residue types could be partly identified in the scalar CCH 3D. Namely, one Glu, Ser, Gln, Leu and two additional Arg **(Fig. 4a).** Comparing these residues with the MAP7 MTBD sequence showed that they are present at the extreme N- or C-terminus. Furthermore, we compared the overall abundance of non-overlapping resonance types to the number of resonances of that type present in the aforementioned terminal domains, suggesting that the experimental signal intensity observed in our scalar based experiments is in reasonable agreement with the relative occurrence of amino-acids in the protein segments Pro 59-Ala 70 and Glu 148-Pro 170 **(Fig. 5b, Supplementary Fig. 4b).** The residue abundance was calculated by dividing the integrated intensities of a resonance region by the average of integrated signal of separated peaks corresponding to one residue. For the scalar experiments, this average was taken from non-overlapping Leu HDCD, Leu HBCB, Arg HDCD and Ile HG2CG2 resonances, and for the dipolar experiment, the average was taken from non-overlapping Ser HBCB, Val CGHG, Leu CBHB and Ala CAHA resonances. Note that our spectral analysis estimates a larger fraction of flexible Leu side chains than present in the N- and C- terminal region, which would be compatible with sidechain motion even in the helical binding region. For example, Leu 103 is located between tubulins and might therefore exhibit flexibility.

#### Analysis of putative MAP7 MTBD A83-V87 beta-strand

By verifying connections in the CAHA region of the dipolar spectrum corresponding to the aforementioned β-strand resonances, we could identify Ala, Glu, Arg and a CAHA correlation corresponding to Val, Ile or Thr **(Supplementary Fig. 4b).** The presence of an Ala is supported by the peak integral of the residue in the CCH spectra which exceeds the expected number of Ala in the MAP7 sequence. **(Fig. 5b and 5c).** The same is the case for Val. Therefore, the CAHA correlation attributable to Val, Ile or Thr mentioned above might be a Val **(Supplementary Fig. 4b, right)**. Additionally, the resonance could be attributed to Ile 86 because there are two resonances observed for Ile 86 CB-HB and CG-HG in the dipolar 2D spectra, even though the MAP7 MTBD sequence contains only one Ile **(Fig. 4b).** Hence, this correlation remains unidentified. The residues that we identified for the β-strand resonances agree with residues Ala 83, Arg 84, Glu 85, Ile 86 and Val 87. These residues would be in line with the aggregation propensity for MAP7 evaluated by AGGRESCAN, which claims residues 84-89 to be aggregation-prone [1]. Interestingly this region corresponds to the hinge, that was observed in the free MAP7 MTBD α-helix (residue 84-87) [2]. The rest of the aggregate could not be observed in the NMR spectra. This might be due to great heterogeneity or intermediate exchange dynamics.
